# NeoCOMM: A Neocortical Neuroinspired Computational Model for the Reconstruction and Simulation of Epileptiform Events

**DOI:** 10.1101/2024.01.04.574141

**Authors:** Mariam Al Harrach, Maxime Yochum, Giulio Ruffini, Fabrice Bartolomei, Pascal Benquet, Fabrice Wendling

## Abstract

Understanding the pathophysiological dynamics which underline interictal epileptiform events (IEEs) such as epileptic spikes, spike-and-waves or High-frequency oscillations (HFOs) is of major importance in the context of neocortical refractory epilepsy, as it paves the way for the development of novel therapies. Typically, these events are detected in local field potential (LFP) recordings obtained through depth electrodes during pre-surgical investigations. Although essential, the underlying pathophysiological mechanisms for the generation of these epileptic neuromarkers remain unclear. The aim of this paper is to propose a novel neurophysiologically relevant reconstruction of the neocortical microcircuitry in the context of epilepsy. This reconstruction intends to facilitate the analysis of a comprehensive set of parameters encompassing physiological, morphological, and biophysical aspects that directly impact the generation and recording of different IEEs. Accordingly, a novel microscale computational model of an epileptic neocortical column was introduced. This model incorporates the intricate multilayered structure of the cortex and allows for the simulation of realistic interictal epileptic signals. The proposed model was validated through comparisons with real IEEs recorded using intracranial stereo-electroencephalography (SEEG) signals from both humans and animals. Using the model, the user can recreate epileptiform patterns observed in different species (human, rodent, and mouse) and study the intracellular activity associated with these patterns. Our model allowed us to unravel the relationship between glutamatergic and GABAergic synaptic transmission of the epileptic neural network and the type of generated IEE. Moreover, sensitivity analyses allowed for the exploration of the pathophysiological parameters responsible for the transitions between these events. Finally, the presented modeling framework also provides an Electrode Tissue Model (ETI) that adds realism to the simulated signals and offers the possibility of studying their sensitivity to the electrode characteristics. The model (NeoCoMM) presented in this work can be of great use in different applications since it offers an *in silico* framework for sensitivity analysis and hypothesis testing. It can also be used as a starting point for more complex studies.

## Introduction

Epilepsy is defined as a chronic neurological disease that is considered an important cause of disability and mortality ***Beghi (2020)***. It is characterized by spontaneous recurrent seizures which affect all age ranges and genders. Over 70 million people worldwide suffer from epilepsy ***Thijs et al. (2019)***. About one-third of epilepsy patients experience seizures that cannot be effectively managed with anti-epileptic drugs, leading to their classification as pharmacoresistant ***Löscher et al. (2020)***. In such cases, surgery remains a viable option for only a small fraction of patients, typically ranging from 15 % to 20%, who exhibit focal, well-defined, and accessible Epileptogenic Zones (EZ) ***Baud et al. (2018)***. Therefore, the precise delineation of the EZ plays a pivotal role in the success of resection surgery. This delineation typically relies on biomarkers derived from electrophysiological recordings, primarily Local Field Potentials (LFPs) collected via intracortical Stereo-ElectroEncephalography (SEEG) electrodes ***An et al. (2020)***. These biomarkers correspond to epileptiform events observed during ictal (seizures) and interictal (spikes, High-frequency oscillations,…) states. However, although seizures are generally unpredictable and infrequent, Interictal Epileptiform Events (IEEs) are considerably more frequent ***Smith et al. (2022)*** which makes them a valuable and important asset.

Still, the pathophysiological mechanisms underlying the generation of different IEEs and their relationship to ictal activity are still poorly understood ***Aeed et al. (2020)***. In particular, for neocortical focal epilepsies, the complexity of the multilayered structure of the cortex coupled with the altered excitation-inhibition balance induces distinct patterns within intrinsic and diverse neural firing properties. The exact details of these dynamics that allow for the generation of either Interictal Epileptic Spikes (IESs), Interictal Spike and Waves (SWs), or High-frequency Oscillations (HFOs) remain ambiguous ***Aeed et al. (2020)***; ***de Curtis et al. (2012)***. Unveiling their underlying neurobiological mechanisms can offer a better interpretation of the SEEG signals recorded during the pre-surgical diagnostic studies ***Aeed et al. (2020)***.

Over the past decade, research has increasingly emphasized the importance of advancements in computational modeling to enhance the postoperative outcomes of epilepsy surgery ***Rigney et al. (2021)***; ***An et al. (2020)***. These computational methods encompass artifficial intelligence-based and biophysical *in-silico* modeling approaches. Their primary goal is to gain a deeper understanding of the pathophysiological mechanisms that drive the occurrence of epileptic events, with the aim of providing valuable insights into tailored, patient-specific therapeutic approaches ***An et al. (2020)***. In the case of artificial intelligence-based models, they are usually limited by a small number of sample sizes which can decrease their accuracy and predictive efficacies ***Rigney et al. (2021)***. In the context of physiologically relevant models, previous studies have produced intricate and highly complex models of healthy cortical tissue ***Markram et al. (2015)***. Nonetheless, these models require significant computational resources and prove overly intricate for specific types of analyses.

In this study, we present a new neuro-inspired microscale model of the multilayered neocortical column that reproduces the main physiological features of the cortex microcircuitry. This digital reconstruction of the cortical volume incorporates a sufficiently large number of cells considering the diversity of neuron and interneuron types and their electrophysiological firing patterns as well as the complex inter and intracortical connectivity between them. Using the forward modeling scheme, the proposed model is able to simulate realistic LFPs as observed in electrophysiological recording using SEEG electrodes (Figure 11) and in vivo using microelectrodes to record the LFP during epileptogenesis following the iron-chloride mouse model ***Jo et al. (2014)*** (Figure 12). Accordingly, This model was used to explore how distinct intrinsic cell characteristics, when coupled with modified synaptic dynamics and synchronized external inputs, can give rise to particular types of IEEs. It strikes a balance between the complexity of electrophysiological aspects and computational speed.

## Results

### Pathophysiological dynamics of epileptiform events generation

#### Creating an epileptic network

We used the NeoCoMM computational model to create an epileptic tissue that is able to simulate realistic IEEs. This was achieved, first, by adjusting the physiological parameters of the different cells to create a multilayered hyperexcitable network that mimics an epileptic cortical column. Second, the input from the Distant Cortex (DC) was rendered synchronous by decreasing the standard deviation of the stimulation epochs of external Principal Cells (PCs) from different layers. The combination of these two conditions and the adjustment of their respective parameters allowed us to simulate different types of IEEs. This simulation scheme is portrayed in Figure 1. In this section, we investigated the underlying pathophysiological parameters that induce the main types of IEEs that are usually observed in intracerebral EEG recordings and are used as epileptic markers by clinicians. Given that the NeoCoMM computational model permits the simulation of cortical columns in humans, rats, and mice, our investigation will focus on examining the IEEs simulation and generation mechanisms within human and mouse cortical tissues.

**Figure 1.**
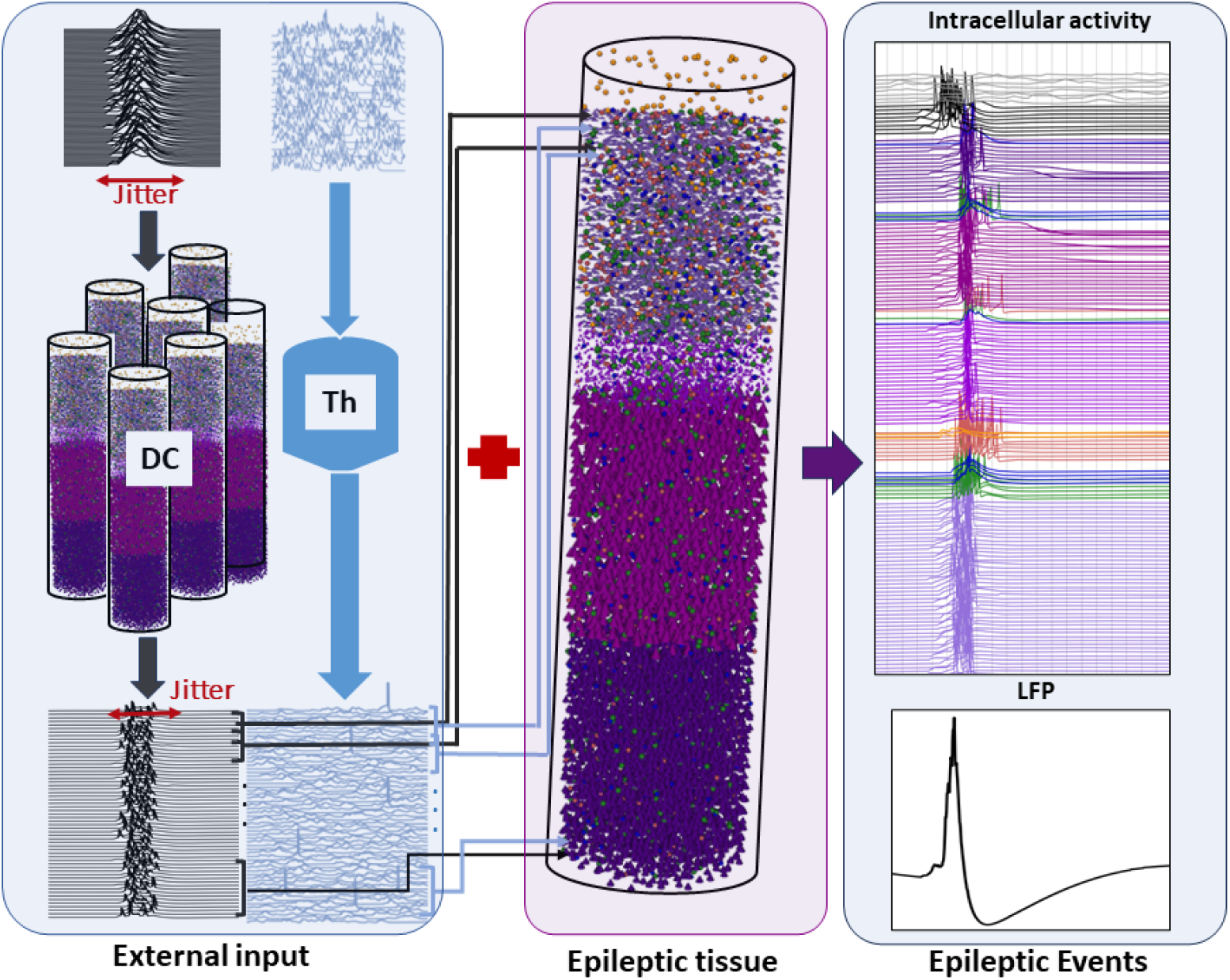
An overview of the Interictal Epileptiform Events (IEEs) simulation scheme. The simulation of IEEs requires a combination of two main elements: An epileptic tissue characterized by pathophysiological hyperexcitable network and a synchronous input of afferent volley of Action Potentials (APs) from the distant cortex (DC). The level of synchrony is described by the stimulation jitter. The neocortical column also receives external input from the Thalamus (Th). The epileptic event simulated in this figure depicts an Interictal Epileptic Spike (IES). The modified synaptic parameters adjusted to obtain this activity are as follows: for PYR cells the *g*_*AMPA*_, *g*_*NMDA*_, and *g*_*GABA*_ were set to 9.61, 0.47, and 36 *mS*/*cm*^2^ respectively.

#### Generation of interictal events in the human neocortical tissue

Starting from the default configuration of the computational model (See section Neocortical Computational Microscale Model (NeoCoMM)), we studied the network activity in response to an external volley of APs coming from the PCs of the DC and induced by a quasi-synchronous stimulus obtained by adjusting the number of individual stimulating inputs and their jitter (their standard deviation). With the default physiological values of the neocortical tissue, no response or very low activity was observed in the recorded LFP which highlights the necessity of both conditions previously mentioned to simulate IEEs.

Accordingly, to simulate different IEE patterns including Interictal Spikes (IESs), Spike Waves (SWs), Double Spikes (DSs) and waves (DSWs), and High-frequency Oscillations (HFOs) (ripples, and Fast Ripples (FRs)), we conducted a simulation study of the parameters involved in the generation of these events. This investigation consisted mainly of studying the impact of pathophysiological synaptic transmission between GABAergic and Glutamatergic cells in the different layers of the neocortical column along with the impact of the external DC input synchrony and intensity. In detail, we focused on the conductances of excitatory (*AMP A*_*R*_ and *NMDA*_*R*_) and inhibitory (*GABA*_*R*_) synaptic receptors as well as on the reversal potentials of GABAergic postsynaptic currents simultaneously with the jitter value and the number of afferent APs (Figure 1).

Figure 2 shows the simulation results for SW, DSW, ripple, and FR compared to real clinical IEEs as depicted in both time and frequency domains. It demonstrates the ability of our model to efficiently reproduce these different interictal patterns by adjusting the underlying dynamics of the epileptic network. For the SW generation (Figure 2.A), simulations indicated that asynchronous input (higher jitter) is needed for the external stimulation of the network along with an increase in the excitatory conductances (*AMP A*_*R*_ and *NMDA*_*R*_) and the GABA reversal potential of postsynaptic current generated at the soma and dendrites of PCs in all layers. In the case of DSW (Figure 2.B) same conditions were applied on the cortical column as for the SW except for the jitter of the input from DC which was reduced (5 *ms* for DSW instead of 8 *ms* for SW). Figures 2.C and 2.D depict real (left) versus simulated (right) Ripples and FR respectively. The ripples whether in the real or simulated signals are characterized by a signal power in a frequency band of 80 to 200 *Hz* as opposed to FR that have a signal power between 200 and 600 *Hz*. To simulate HFOs, the NMDA postsynaptic current conductance of PYR cells was moderately increased to create a hyperexcitable tissue but not to induce the depolarisation of a high number of PYR cells (Figure 2-figure supplement 3 and Figure 2-figure supplement 4). In addition to this condition, the excitatory input from DC was set to be of higher intensity for the FR simulation compared to the ripples with a jitter value between 4 and 5 *ms*. This resulted in a weakly synchronized firing of a set of PC cells (< 15%) that was higher in the case of ripples (< 45%).

**Figure 2.**
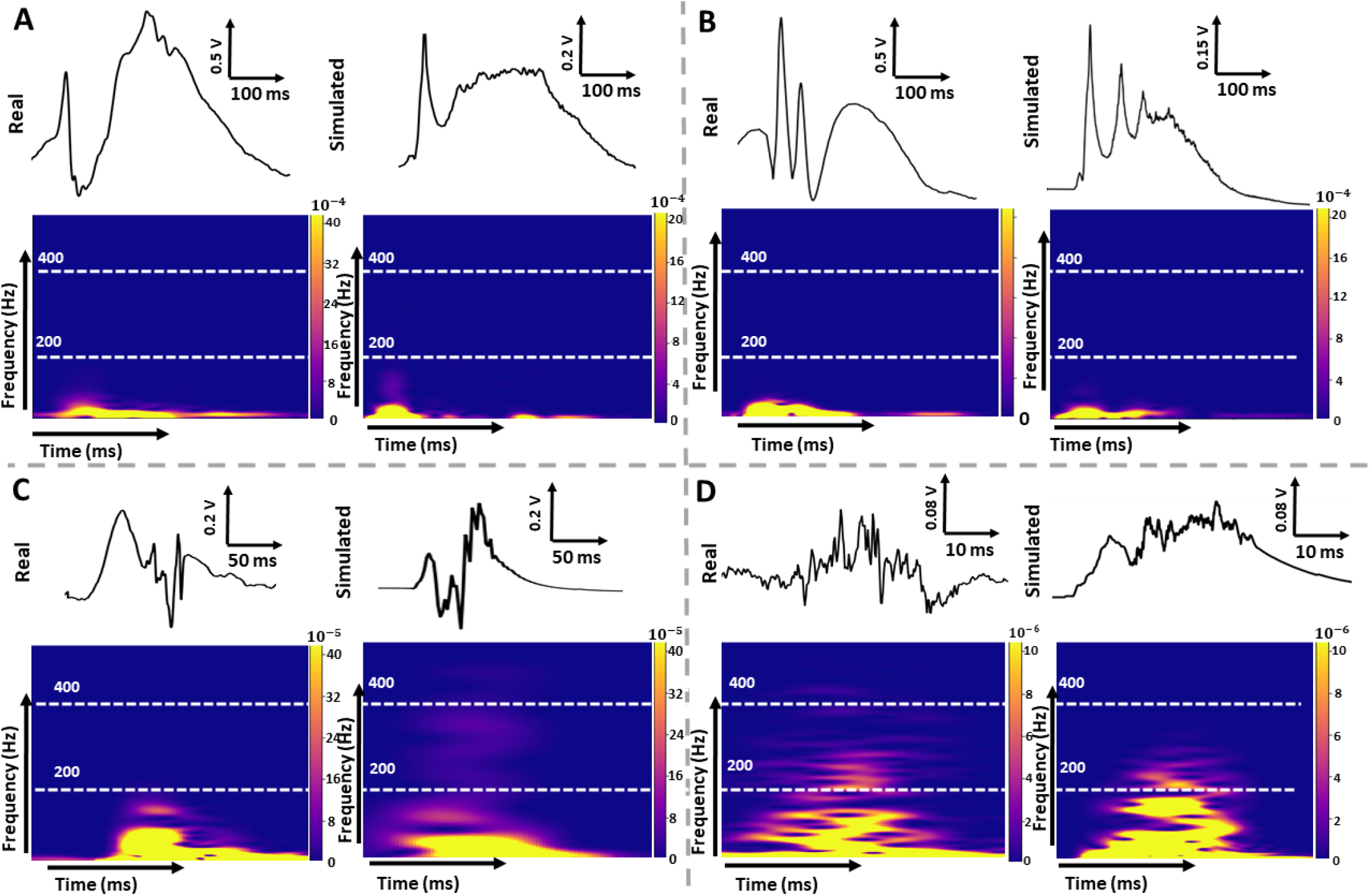
Comparison between clinical and simulated Local field potentials (LFPs) during Interictal Epileptiform Events (IEEs): Spike-and-Wave (SW) (A), Double Spike-and-Wave (DSW) (B), High-Frequency Oscillations (HFOs) (ripples (C) and Fast Ripples (FRs) (D)). Real LFPs were obtained from an epileptic patient with neocortical Temporal Lobe Epilepsy (TLE) using depth SEEG electrodes and portray typical IEE events in time (up) with the corresponding spectrogram (bottom). A: Simulated SW (right) with a rapid component with high amplitude (spike) and a slow wave reflected in the frequency spectrum portraying key elements of the clinically recorded SW (left). This SW was obtained using the following synaptic adjustments: *g*_*AMP A*_ = 9, *g*_*NMDA*_ = 0.67,*g*_*GABA*_ = 31,*E*_*GABA*_ = −67 and input Jitter =8 *ms*. B: Simulated DSW (right) with two successive rapid components with high amplitudes and a slow wave. The corresponding synaptic parameters are as follows: *g*_*AMP A*_ = 7.61, *g*_*NMDA*_ = 0.80,*g*_*GABA*_ = 25,*E*_*GABA*_ = −73 and input Jitter =5 *ms*, C: Simulated (Right) and real (left) LFPs with interictal R characterized by a frequency range between 80 and 200 *Hz*. The adjusted parameters for this simulation are: *g*_*AMP A*_ = 7.38, *g*_*NMDA*_ = 0.48,*g*_*GABA*_ = 38,*E*_*GABA*_ = −74, D: Simulated (Right) and real (left) LFPs with interictal FR characterized by a frequency range between 200 and 600 *Hz*. The following parameters values were used to obtain this simulation: *g*_*AMP A*_ = 7.6, *g*_*NMDA*_ = 0.5,*g*_*GABA*_ = 37,*E*_*GABA*_ = −74 and input Jitter =5 *ms* **Figure 2—source code 1.** The configuration files for the simulations in NeoCoMM: https://gitlab.univ-rennes1.fr/myochum/neocomm/ **Figure 2—figure supplement 1.** The intracellular activity corresponding to the SW signal in (A) **Figure 2—figure supplement 2.** The intracellular activity corresponding to the DSW signal in (B) **Figure 2—figure supplement 3.** The intracellular activity corresponding to the HFOs signal in (C) **Figure 2—figure supplement 4.** The intracellular activity corresponding to the FR signal in (D)

Another advantage of the model is its ability to simultaneously display the extracellular (Figure 3.A) and intracellular (Figure 3.B) activity of all the network cells. This highlights the capability of the model to uncover the specific underlying mechanisms responsible for generating each type of epileptiform pattern. Comparing the intracellular response with the corresponding LFP signal and analyzing the adjusted electrophysiological parameters used to obtain the IEE pattern, allowed us to elucidate and examine these distinct mechanisms. In this regard, Figures 3 and 4 presents IES and SW simulations, respectively, compared to real signals, along with the corresponding intracellular activity of PCs and interneurons in the five neocortical layers. In the case of IES (Figure 3), it is characterized by a sharp wave lasting between 50 and 100 *ms* followed by a brief negative wave (< 50*ms*). By examining the corresponding simulated intracellular activity (Figure 3.B), we observed that during the sharp spike component, all PC cells in the column were depolarized and exhibited high synchrony, with some cells firing several APs within a brief individual time frame (< 30 *ms*). Similarly, interneuron activity was synchronous. However, the discharge period extended beyond that of the PCs, resulting in the negative wave following the spike. To obtain the simulation presented in Figure 3, the default parameters of the model were adjusted by increasing the conductances *AMP A*_*R*_ and *NMDA*_*R*_ of PCs and increasing the reversal potential of GABAergic postsynaptic currents. The jitter value of the external input from DC was set to 4 *ms*.

**Figure 3.**
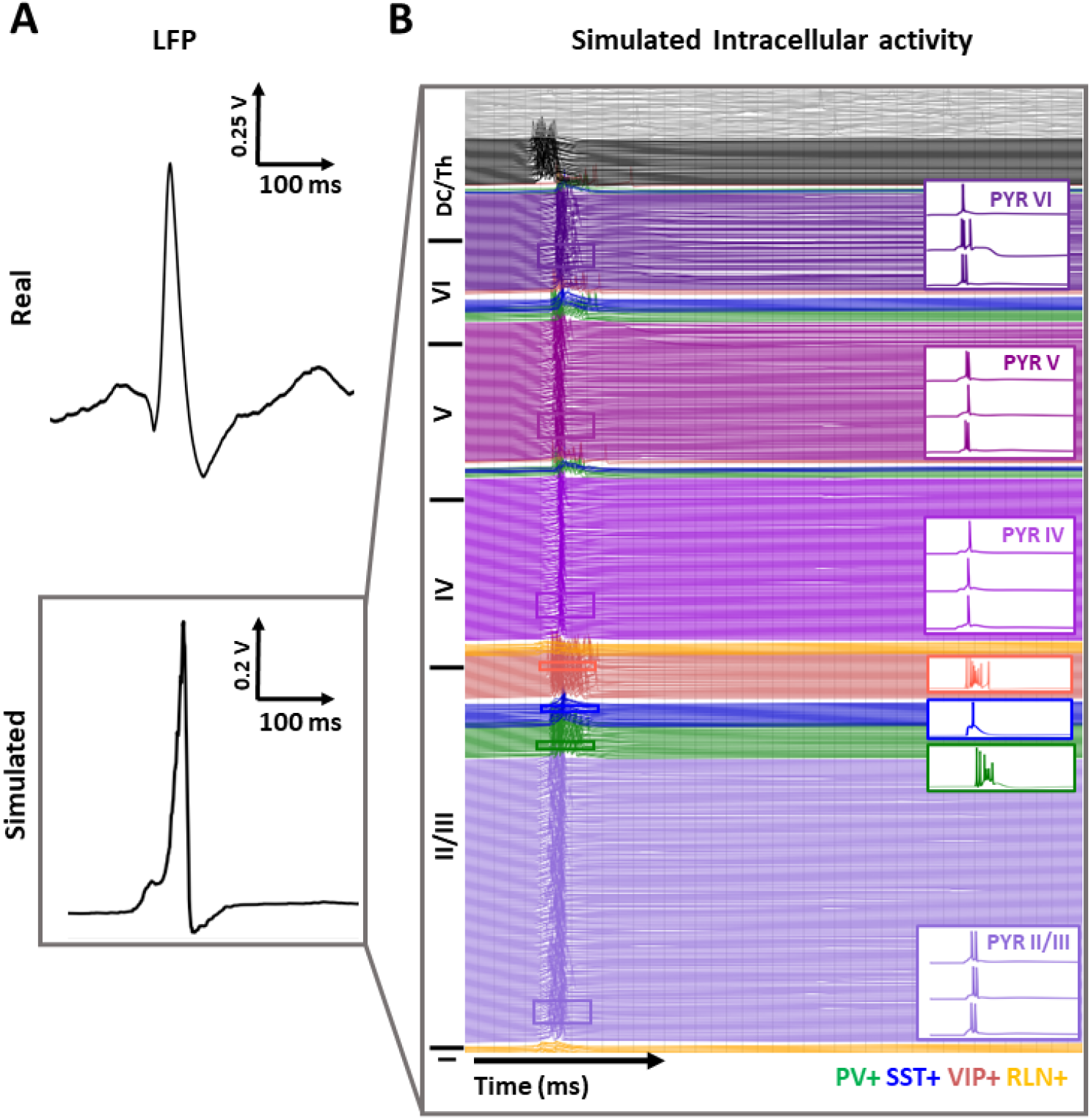
Interictal Epileptiform Spikes (IESs) simulation with the NeoCoMM model. A: Comparison between real and simulated local field potential (LFP) during an IES. (A, top) Typical clinical IES recorded with depth EEG electrodes in the transverse temporal gyri of Heschel from a patient with neocortical temporal lobe epilepsy. (A, bottom) Simulated IES generated by the computational model NEOCOMM. (B) Overview of the intracellular activity corresponding to the simulated LFP in (A, bottom). 30% of cellular activity is shown for the different cell types: Pyramidal cells (PYR), Parvalbumin expressing interneurons (PV+), Somatostatin expressing interneurons (SST+), vasoactive intestinal polypeptide expressing interneurons (VIP+) and Reelin expressing interneurons (RLN +). The synaptic parameters of the PYR cells were adjusted from default values in order to create an epileptic tissue. These values are: *g*_*AMP A*_ = 8.76, *g*_*NMDA*_ = 0.63,*g*_*GABA*_ = 37,*E*_*GABA*_ = −66. The input Jitter was set to 4 *ms* **Figure 3—source code 1.** The configuration file for the simulation in NeoCoMM: https://gitlab.univ-rennes1.fr/myochum/neocomm/

Following the same approach, we simulated the SW presented in Figure 4. We noticed that the model successfully mirrored the key elements of the recorded SW (Figure 4.A), both in time and frequency content characteristics. Concerning the physiological parameters adjusted to obtain this simulation compared to the IES in Figure 3, the DC stimulation jitter was increased (5 *ms* instead of 4 *ms*) as well as the intensity and number of stimulation epochs resulting in a higher afferent volley from the DC to the epileptic column. Additionally, along with increasing the glutamatergic conductances of synaptic receptors, we decreased the GABAergic conductance of synaptic receptors for all PC cells. Upon analyzing the intracellular activity responsible for generating the SW pattern (Figure 4.B), we observed that, in the same manner as the IES, the spike component resulted from highly synchronous APs from PCs. However, in this case, the discharges from PCs didn’t completely cease for all PCs after about 30-40 *ms*; instead, they continued for some time, triggering a new wave of depolarization in all network cells (Figure 4.B). For the second wave, the APs were highly asynchronous and took longer to dissipate (>150 *ms*). These asynchronous bursts of APs determined the wave’s shape, including its duration and amplitude. For example, the simulated SW in Figure 2.A exhibited a wave with a much longer duration and higher amplitude. This was a result of a more pronounced asynchronous bursting in all cells, as shown in Figure 2-supplement figure 1. Compared to the SW of Figure 4, this difference can be attributed to higher *AMP A*_*R*_ and GABAergic reversal potential for postsynaptic currents in PC cells.

**Figure 4.**
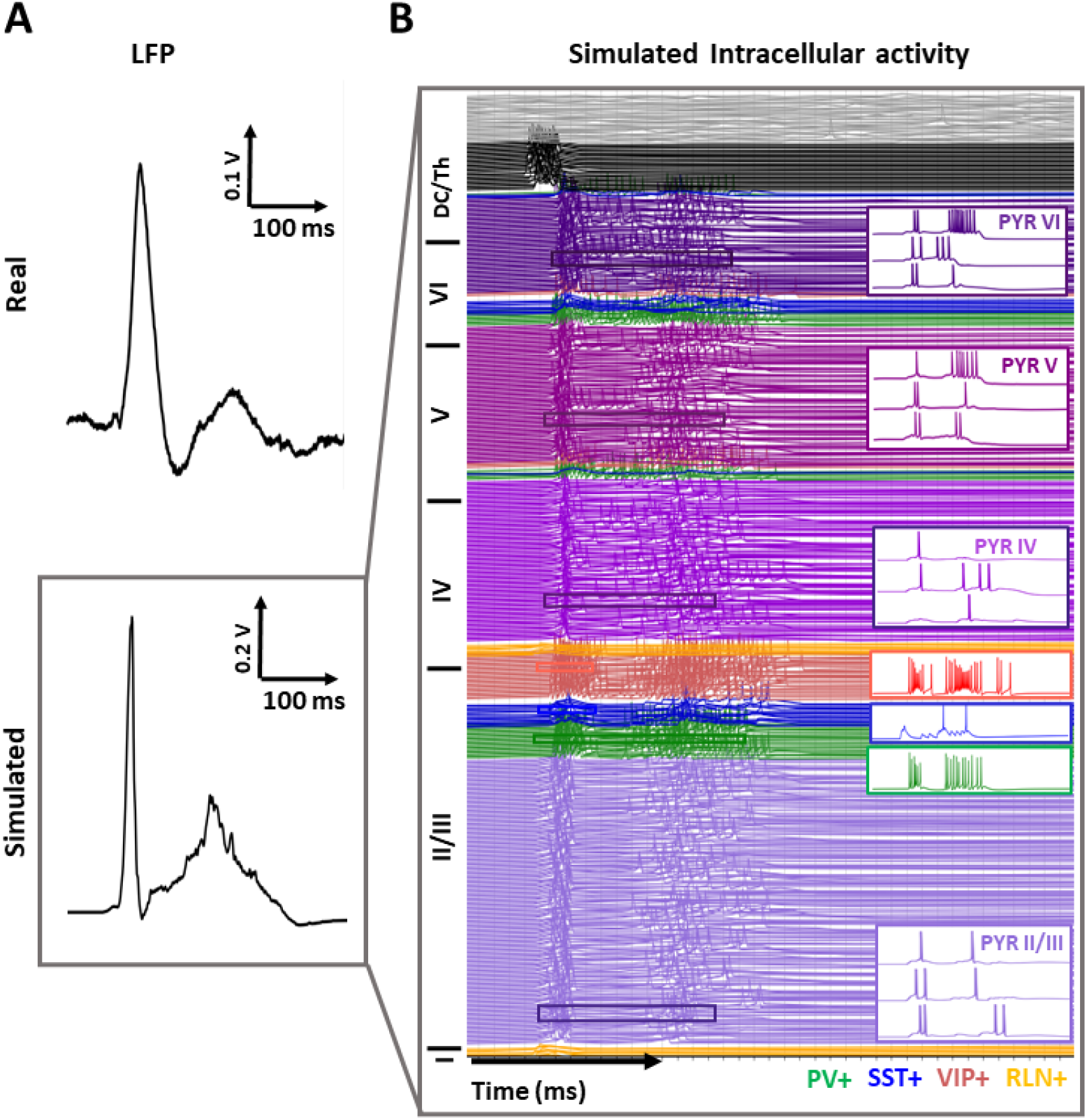
Interictal Epileptiform Spikes and Wave (SW) simulation with the NeoCOMM model. A: Comparison between real and simulated local field potential (LFP) during an SW. (A, top) Typical clinical SW recorded with depth EEG electrodes in the transverse temporal gyri of Heschel from a patient with neocortical temporal lobe epilepsy. (A, bottom) Simulated IES generated by the computational model NEOCOMM. (B) Overview of the intracellular activity corresponding to the simulated LFP in (A, bottom). 30% of cellular activity is shown for the different cell types: Pyramidal cells (PYR), Parvalbumin expressing interneurons (PV+), Somatostatin expressing interneurons (SST+), vasoactive intestinal polypeptide expressing interneurons (VIP+) and Reelin expressing interneurons (RLN +). The synaptic parameters of the PYR cells were adjusted from default values in order to create an epileptic tissue. These values are as follows: *g*_*AMP A*_ = 7.25, *g*_*NMDA*_ = 0.65,*g*_*GABA*_ = 21,*E*_*GABA*_ = −72. The input Jitter was set to 6 *ms* **Figure 4—source code 1.** The configuration file for the simulation in NeoCoMM: https://gitlab.univ-rennes1.fr/myochum/neocomm/

To assess the impact of the electrophysiological model parameters on the type and morphological features of the simulated IEEs, we conducted a sensitivity analysis centering around the excitatory and inhibitory synaptic parameters. For each studied parameter and each IEE type, all other model parameters were fixed during the simulations. We started by studying the impact of the DC input synchrony level on the peak spike amplitude and duration for IESs, SWs, and DSWs. Figure 5.A showed that the level of synchronization of the external cells’ firing pattern has a direct influence on the morphology of the simulated IEE. In the case of IES, reduced input jitter value (below 4 *ms*) resulted in a very low amplitude signal (Figure 5.A). Above 4 *ms* the spike amplitude decreased and its duration increased with input synchronization decrease (jitter increase). Similarly, in the case of SW and DSW, the amplitude of the spike peak was higher for higher external APs synchrony. Their duration, however, seemed to increase for the SW and decreased for the DSW with the jitter value increase.

**Figure 5.**
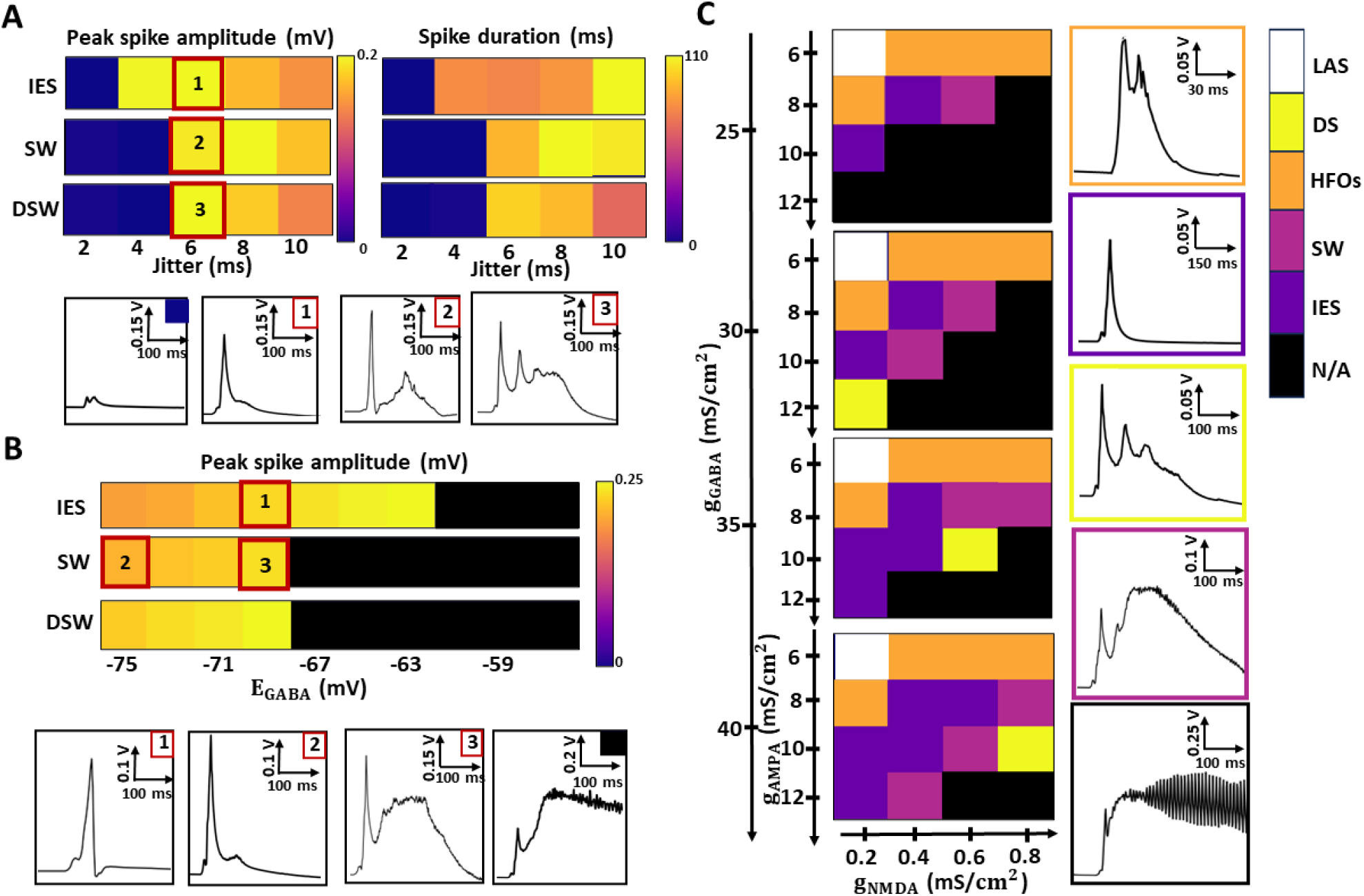
Sensitivity analysis of simulated Interictal Epileptiform Events (IEEs) types and morphological features with model parameters (electrophysiological synaptic parameters). (A) Impact of the input Jitter of the peak spike amplitude and duration of Interictal Epileptic Spikes (IESs) and Spike and Waves (SWs). (B). Impact of the GABA reversal potential value (*E*_*GABA*_) on the peak spike amplitude for IESs and SWs. (c) Color-coded maps illustrating the impact of *AMP A*_*R*_, *NMDA*_*R*_, and *GABA*_*R*_ conductances on the type of simulated IEEs. LAS: Low Amplitude Signal, DSW: Double Spike and Wave, HFOs: High Frequency Oscillations, SW: Spike-and-Wave, IES: Interictal Epileptic Spike, N/A: oscillatory signal similar to ictal activity.

For these three IEEs (IESs, SWs, and DSWs), we also studied the impact of the GABA reversal potential *E*_*GABA*_ of postsynaptic GABAergic currents on the simulated events shape. As depicted in Figure 5.B, the spike component amplitude was higher for higher *E*_*GABA*_ values. For very hyperexcitable neural networks, we obtained an oscillatory activity that appears like ictal activity and is characterized by continuous bursting of all the network’s cells (black squares in Figure 5.B). The threshold for this ictal activity was found to be higher for IESs compared to SWs and DSWs. Another interesting finding was that for the SW, the simulated events switch from IES to SW between -75 and -69 *mV* as shown in the IEEs plots in Figure 5.B red squares 2 and 3.

Finally, we focused on the postsynaptic conductance values associated with *AMP A*_*R*_, *NMDA*_*R*_, and *GABA*_*R*_ receptors of PCs and their impact on the type of simulated IEE. After freezing all other parameters of the model, we simulated the response of the neocortical column to an external stimulation for each triplet conductance value (*g*_*AMP A*_, *g*_*NMDA*_, and *g*_*GABA*_) over predefined intervals. Figure 5.C presents the color-coded maps indicating the different parameter configurations and the corresponding IEE. These configurations are divided into four color maps for each *GABA*_*R*_ conductance value. Then, each colormap portrays the corresponding events of each excitatory pair of *NMDA*_*R*_ and *AMP A*_*R*_ conductances. The boundaries of these conductances were chosen by keeping in mind physiological realism. Our analysis revealed a repetitive pattern in the case of the low amplitude signal and HFOs squares. This implies that independently from the inhibitory postsynaptic current intensity of PCs, very low excitatory postsynaptic currents (*g*_*AMP A*_ = 6*mS*∕*cm*^2^ and *g*_*NMDA*_ = 0.2*mS*∕*cm*^2^) cannot simulate epileptic signals resulting in a low amplitude signal or physiological LFP. Also, independently from the inhibitory conductance value of PC’s postsynaptic receptors, setting *g*_*AMP A*_ to 8 *mS*∕*cm*^2^ or increasing *NMDA*_*R*_ conductance (*g*_*NMDA*_ > 0.2*mS*∕*cm*^2^) while maintaining the *AMP A*_*R*_ conductance to 8 *mS*∕*cm*^2^, leads to the simulation of HFOs. In the same vein, values (*g*_*AMP A*_, *g*_*NMDA*_) =(10,0.2) and (8,0.4) *mS*∕*cm*^2^ resulted in IES each time. In contrast, for some (*g*_*AMP A*_, *g*_*NMDA*_) values the simulated event switched from IES to SW with increasing inhibitory synaptic conductance *g*_*GABA*_. This implies that the neocortical tissue needs to be sufficiently hyperexcitable with increased excitatory input currents and inhibitory activity to simulate IESs and SWs. Additionally, we need to point out the GABAergic postsynaptic current importance wherein its decrease results in a much more hyperexcitable network leading to an oscillatory activity that resembled ictal discharges (plot in black square in Figure 5.C). In the case of DSW, specific conductance combinations are needed, these triplet values are (*g*_*AMP A*_, *g*_*NMDA*_, *g*_*GABA*_) = (10,0.2, 25), (10,0.6,30), (12,0.2,35) *mS*∕*cm*^2^. The sensitivity analysis presented allowed us, not only, to investigate the impact of certain parameters on the simulated IEEs but also provided a guideline for simulations with NeoCOMM. A summary of the pathophysiological parameter values for the simulation of different interictal epileptic patterns is provided in Appendix 1-Table 1.

#### Generation of interictal events in the mouse cortical tissue

As mentioned earlier, the NeoCoMM model can simulate the cortical column and microscopic activity in humans, rats, or mice. In this section, we investigated the underlying mechanisms for creating epileptic networks that can generate IEEs in the mouse’s cortical tissue. After performing a wide range of simulations, we found that in order to simulate IES, we need a combination of parameter configurations: Firstly, the external input from the DC needs to be highly synchronous. Secondly, and similarly to the human case, the cortical column needs to be hyperexcitable. However, to simulate this hyperexcitability, we needed to adjust the synaptic parameters of both PCs and interneurons. Specifically, for the simulation of an IES, the following adjustments to the synaptic currents are required in all layers of the simulated neocortical volume: 1) an increase in excitatory conductances (*AMP A*_*R*_ and *NMDA*_*R*_) for PC cells, 2) an increase in *AMP A*_*R*_ conductances for PV, SST, and VIP cells. The combination of these criteria creates an epileptic network that can generate IES events in response to a volley of synchronized APs from the DC. An example of a simulated IES is presented in Figure 6.A. As depicted, with the appropriate parameter settings, the model effectively replicated an IES, demonstrating its reliability when compared to real *in vivo* recording.

**Figure 6.**
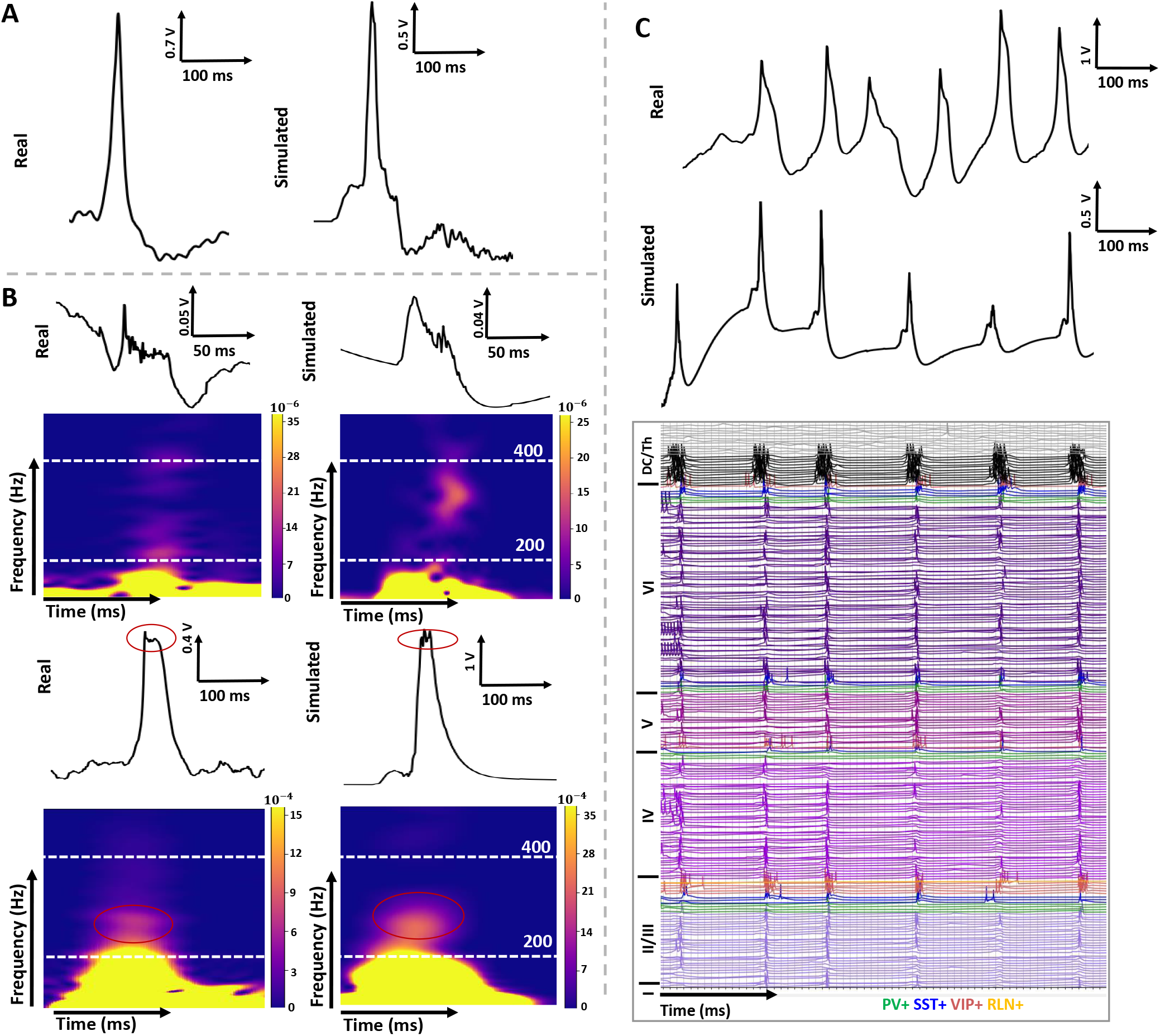
Comparison between experimental and simulated neocortical Interictal Epileptiform Events (IEEs) in mice. (A) Real vs. simulated Interictal Epileptic Spike (IES). The following synaptic adjustments were made to obtain this simulation: *P C*∕*g*_*AMP A*_ = 12, *P C*∕*g*_*NMDA*_ = 0.6, *P V* ∕*g*_*AMP A*_ = 12, *SST* ∕*g*_*AMP A*_ = 8, *SST* ∕*g*_*GABA*_ = 2, and *V IP* ∕*g*_*AMP A*_ = 8 *mS*∕*cm*^2^. (B) Examples of two different types of simulated FRs compared to recorded *in Vivo* ones. (up) A standalone FR obtained for *P C*∕*g*_*AMP A*_ = 9.7, *P C*∕*g*_*NMDA*_ = 0.6, *P V* ∕*g*_*AMP A*_ = 6, *P V* ∕*g*_*GABA*_ = 2, *SST* ∕*g*_*AMP A*_ = 8, *SST* ∕*g*_*GABA*_ = 2, *V IP* ∕*g*_*AMP A*_ = 8 *mS*∕*cm*^2^ and *J itter* = 6*ms*. (bottom) A FR superimposed on a spike obtained for *P C*∕*g*_*AMP A*_ = 10, *P C*∕*g*_*NMDA*_ = 0.6, *P V* ∕*g*_*AMP A*_ = 4, *P V* ∕*g*_*GABA*_ = 4, *SST* ∕*g*_*AMP A*_ = 8, *SST* ∕*g*_*GABA*_ = 2, *V IP* ∕*g*_*AMP A*_ = 8 *mS*∕*cm*^2^. (C) Simulation of repetitive epileptic discharges (up) with the corresponding intracellular activity. This simulation was obtained for the same settings as in (A) with a periodic external input of 8 Hz. Outside the indicated parameter values that have been used for these specific simulations, the default parameter values defined in the NeoCOMM model were employed. The in vivo recordings were obtained from an epileptic mouse following the iron ion model described in section. **Figure 6—source code 1.** The configuration files for the simulations are provided in: https://gitlab.univ-rennes1.fr/myochum/neocomm/ **Figure 6—figure supplement 1.** The intracellular activity corresponding to the IES signal in (A) **Figure 6—figure supplement 2.** The intracellular activity corresponding to the FR type 1 signal in (B) **Figure 6—figure supplement 3.** The intracellular activity corresponding to the FR type 2 signal in (B)

For the simulation of HFOs and particularly FRs, we used the model to uncover the mechanisms responsible for the generation of different types of FR that are usually seen in experimental recordings in mice. In this regard, we chose the two widespread types of FR; isolated High-frequency oscillations in the FR frequency band (200-600 *Hz*) and FR segment superimposed on a spike component. An example of these two types is depicted in Figure 6.B. Going forward we will refer to the isolated FR type as FR type 1 (Figure 6.B, up) and to the one cooccuring with a spike as FR type 2 (Figure 6.B, bottom). According to the simulations, and compared to IESs, FRs are the result of an asynchronous firing of a small number of PCs throughout the cortical layers (Please refer to Figure 6. Figure supplement 2 and 3). A closer analysis indicated that for FR type 1, all PYR cells were depolarised, but only a small percentage attained the threshold to fire out-of-phase APs (< 10%). For the same external input as IES, FR Type 1 was obtained by reducing the *AMP A*_*R*_ conductance of PC cells from 12 *mS*∕*cm*^2^ (for IES) to 8 *mS*∕*cm*^2^ and by increasing the *GABA*_*R*_ conductance of PV+ interneurons from 1.38 to 2 *mS*∕*cm*^2^. These Adjustments created a less hyperexcitable network of neurons that fires less and more asynchronously. Interestingly, to simulate the type 2 FR starting with the same parameter configuration as type 1 FR, only the PV+ postsynaptic current parameters needed to be adjusted. These adjustments consisted of decreasing the *AMP A*_*R*_ glutamate conductance to 4 *mS*∕*cm*^2^ and increasing the *GABA*_*R*_ conductance to 4 *mS*∕*cm*^2^. The new physiological synaptic parameters created a new network activity with increased asynchrony with some cells exhibiting a bursting activity (Figure 6-figure supplement 3).

Lastly, to mimic real activity with groups of repetitive IES, we simulated 1 s of activity wherein the epileptic tissue (same configuration as for the IES in Figure 6.A) received external inputs (Volleys of APs) of 8 Hz. These inputs are obtained by applying the same stimulation mechanism described in the methods section (*jitter* = 6*ms*) in a continuous manner. The stimulation epochs were determined based on the 8 Hz stimulation frequency with a random uniform shift of 60*ms* to add realism. The simulation result is shown in Figure 6.C along with an experimental recording showcasing the same type of activity. Comparing the simulated signal to the experimental one highlighted the performance of our model and its ability to not only accurately reproduce interictal patterns but also its ability to offer an insight into the microscopic activity of cells responsible for these patterns. The intracellular responses of individual cells suggested that the peak amplitude of IES is directly linked to the number of synchronous APs fired by PCs. Moreover, the duration between external stimulations was also found to play a role in the shape of the IES. For example, for very brief interspike intervals, some interneurons continue to fire inhibiting the responses for the next spike which results in a lower amplitude IES. This phenomenon is depicted in Figure 6.C in the second to last IES of the simulated signal.

### Impact of the recording electrode on the characteristics of epileptiform events

The NeoCOMM model incorporates a biophysical model of the recording electrode. This model represents both the geometrical properties of the electrode (shape, radius, position, and insertion angle), and the Electrode Tissue Interface (ETI) modeled as an equivalent circuit. In the case of human simulations, we modeled the SEEG electrode that is usually used in clinical settings. A diagram of this electrode is shown in Figure 7.A. For the clinical signals presented in this work, the SEEG electrode used is a typical SEEG electrode with cylindrical Platinum (Pt) contacts of 2 *mm* Heights (*H*) and 0.8 *mm* radii. On the other hand, in the case of mouse recordings, a wire electrode was used with a Stainless Steel (SS) disk contact of 62.5 *μm* radius. The corresponding simulated model is portrayed in Figure 7.B. For both humans and mice, the ETI model was integrated into the simulations of the IEEs with the default values corresponding to each electrode illustrated in Figure 7.

**Figure 7.**
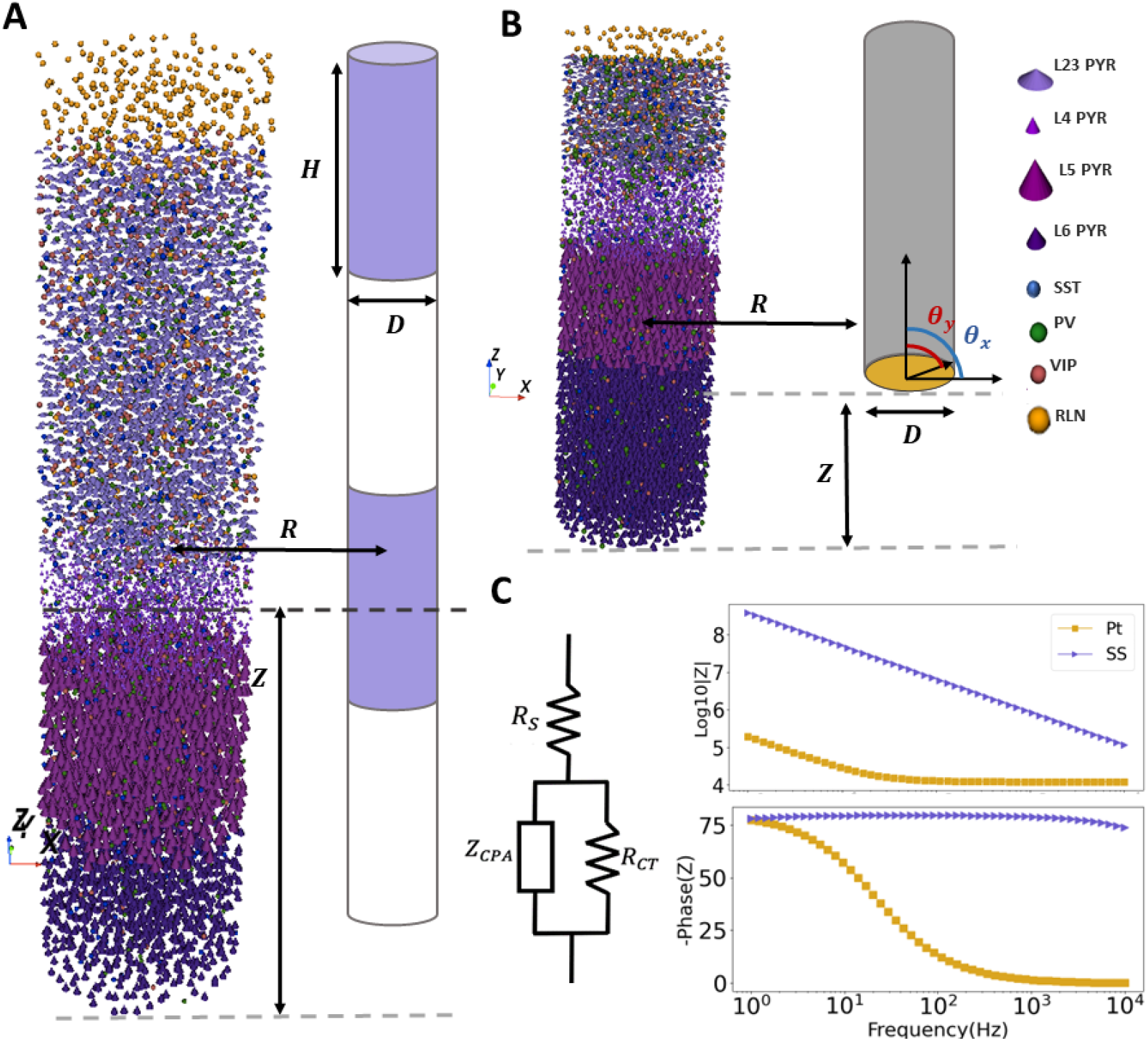
SEEG Recording electrodes simulation. (A) Schematic of the electrodes used in clinical SEEG recordings and simulated for the human neocortical LFP recording. (B) Diagram of the electrode used *in vivo* recordings and simulated for the recording of LFP in mice. (C) The electrode Tissue Interface (ETI) equivalent circuit and corresponding bode plots for Platinum (Pt) electrodes used in clinical settings and stainless steel (SS) used *in Vivo*. The circuit elements consisted of the spreading resistance (*R*_*s*_), the charge transfer resistance (*R*_*CT*_), and the constant phase angle impedance (*Z*_*CP A*_). **Figure 7—figure supplement 1.** Bode plot of the transfer functions for both Pt and SS electrodes

Figure 7.C portrays the impedance variation with frequency for both electrodes. We can notice that for SEEG electrode contacts, the impedance is almost three orders of magnitude lower compared to that of the microelectrode electrode used in mouse recordings. This implies that the blurring effect due to the ETI is more pronounced in the case of microelectrodes compared to SEEG electrodes. However, we have to consider the spatial averaging that a larger recording surface may entail. Moreover, while microelectrodes offer spatial selectivity resulting in higher impedance, SEEG electrodes offer a larger recording field with lower impedance. In particular, the filtering effect of the electrode is more clearly portrayed by the ETI transfer function (Figure 7-supplementary figure 1). For the SEEG electrodes, the cut-off frequency of the ETI filter is around 200 Hz meaning it only distorts the high-frequency contents of the LFP. In the case of microelectrodes, the ETI has a blurring effect for all frequency ranges that is more pronounced for higher frequency oscillations.

Using the electrode model, we studied the effect of the geometrical characteristics of the recording electrode on the recorded IEEs. In particular, we chose to study the different electrode characteristics on the amplitude of the simulated SW. Figure 8 portrays the amplitude variation for both spike and wave parts with the Radius of the electrode contact, the electrode depth, its distance from the epileptic column, and the insertion angle.

**Figure 8.**
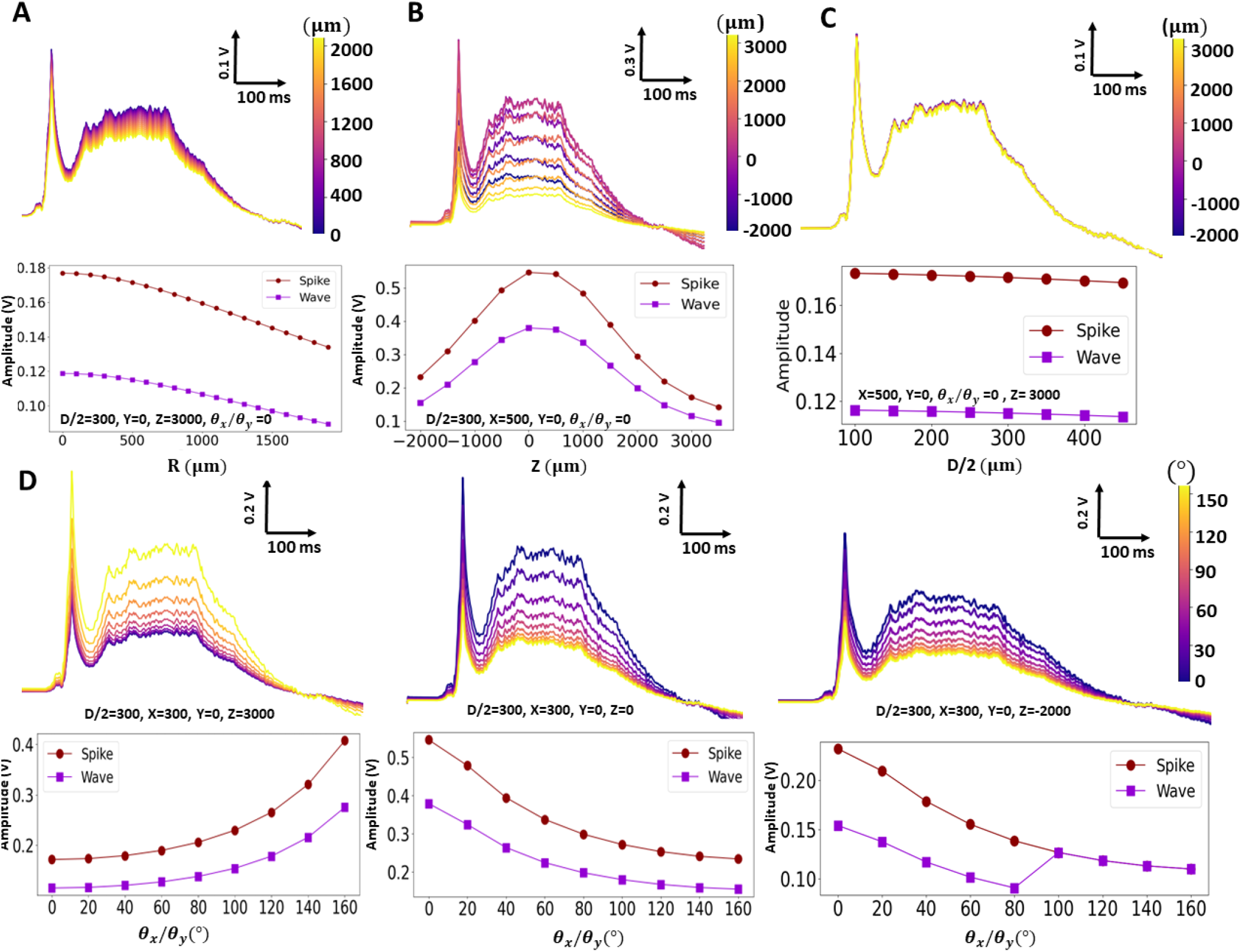
Impact of the SEEG electrode’s geometrical characteristics on the shape of the simulated SW signal for Pt type contacts. (A), (B) and (C) Variation of the spike and the wave amplitudes with the electrode distance (*R*) from the column, depth (*Z*) with respect to the column, and radius (*D*∕2) respectively. (D) Impact of the electrode’s insertion angles (*θ*_*x*_∕*θ*_*y*_) on the spike and the wave amplitudes for *Z* = 3000, 0, and -2000 *μm*. For all the simulations the electrode contact height (*H*) was fixed at 2000 *μm*. All studied parameters are visualized in the electrode’s diagram in Figure 7.

For all the considered parameters both spike and wave components had the same variation profiles. Their amplitude decreased with increasing distance of the electrode contact (Figures 8.A and 8.B). Interestingly, the radius of the electrode contact did not affect the amplitude of the SW (Figures 8.C) which can be explained by the fact that only one column is simulated which is not the case in the cerebral cortex. Lastly, the insertion angle seemed to influence the recorded signal amplitude depending on the position of the electrode’s contact with respect to the cortical column. Accordingly, the amplitudes increased with rotations of the electrode around the x or y axes when the electrode was above the cortical tissue, and decreased when the electrode was below.

## Discussion

This paper presents a novel and physiologically relevant-model of the cortical microcircuitry (NeoCoMM), encompassing realistic neural morphologies, layer dimensions, neural densities, ratios of neuron subtypes, electrophysiology, synaptic physiology, connectivity, and a biophysical recording electrode model. It also provides the possibility to choose from three different species: humans, rats, and mice. This computational model is a simplified but still highly reliable tool to study and uncover mechanisms underlying interictal activity in epilepsy. In this context, it provides the significant ability to simultaneously display both extracellular (local field potentials) and intracellular (action potentials) activity. This capability is especially important in a context where single-cell recordings are challenging to obtain for all involved cell types (excitatory and inhibitory) ***Carlson et al. (2018)***. As a result, it can be used to reveal the behavior of single cells (both PCs and interneurons) during the occurrence of epileptic events, which still remains unclear. This is of major importance since understanding the connection between the response of single cells, their pathophysiological parameters, and the recorded epileptic signal (LFP) can offer new prospects for improved therapeutic options.

In line with these points, in this work, we used NeoCoMM to study the dynamics of interictal epileptiform events (IEEs) in the human neocortex and the mouse cortex. IEES are usually described by physiological and network abnormalities caused by enhanced excitatory connections (hyperexcitable network) or reduced inhibitory connections ***de Curtis et al. (2012)***. In our model, two conditions were combined to simulate IEEs: i) increased synchrony of the DC input and ii) adjustment of parameters involved in excitatory and inhibitory synaptic transmission.

In the case of the human epileptic neocortical column, the model was able to simulate the main types of IEEs including IESs, SWs, DSWs, and HFOs (Ripples and FRs) with high accuracy, as portrayed in Figures 2, 3 and 4. A sensitivity analysis allowed us to investigate the distinct hyperexcitability mechanisms related to each IEE type and their relationship to the pathophysiological parameters of the layered network. In the case of IES, they are usually described as the result of the synchronous firing of a hyperexcitable neural network ***Karoly et al. (2016)***; ***Demont-Guignard et al. (2012)***; ***Lévesque et al. (2018)***. Our simulations found that this could be achieved by creating a hypersynchronous input from the DC (small jitter) that triggers a synchronous firing of single cells (Figure 3) as a result of enhanced glutamatergic postsynaptic potentials at the level of PCs along with increased inhibition threshold. These configurations were consistent with in vivo and in vitro recordings ***Lévesque et al. (2018)*** even though these studies also insist on the heterogeneity of single-cell firing patterns ***Lai et al. (2023)***. We also observed that, depending on the neocortical layer, a bursting phenomenon is portrayed by the PCs which is in line with several studies that pointed out that bursting pyramidal cells play a crucial role in IES initiation in the human neocortex ***Hofer et al. (2022)***; ***Tóth et al. (2018)***.

The results presented also emphasized the critical role of excitatory to inhibitory ratio imbalance on the single-cell dynamics and thence the generated epileptic event. For SW, the model revealed that a slightly decreased synchrony of external input (increased jitter value) combined with an increase of the NMDA conductance, compared to IES network settings, decreases the ability of interneurons to entirely halt the firing of PCs (Figure 4.B). This results in a decreased feedforward inhibition causing the initiation of a second wave of asynchronous firing of PCs due to synaptic transmission governed by interneuron activation. The combination of the first synchronous firing of PCs followed by the volley of asynchronous slower (longer) bursting of cells gives rise to the SW pattern. Another example to underline this mechanism was given in Figure 2.A where an even lower synchronized input was used to trigger the epileptic network characterized by an increased excitability (compared to the previous one) and an increased inhibition threshold of PCs. This example demonstrated the direct impact of these conditions on the amplitude and the duration of the wave (Figure 2-supplementary figure 1). These findings are inconsistent with some studies that claimed that SW patterns are initiated by paroxysmal depolarization shifts ***de Curtis et al. (2012)***; ***Keller et al. (2010)***. However, these same studies also mentioned that the paroxysmal depolarisation hypothesis depended on the brain region and its level of epileptogenicity and presented inconsistent single-unit responses ***de Curtis et al. (2012)***.

Simulations of HFOs (Figure 2.C and 2.D) for the human neocortex confirmed previously obtained findings using computational modeling for the simulation of HFOs ***Demont-Guignard et al. (2012)*** in the hippocampus. These findings suggested that HFOs are the result of the weakly synchronized firing patterns in a small subset of PCs (Figures 2 - figure supplement 3, 4). Nevertheless, due to the multilayered nature of the neocortical tissue and the complex interconnectivity between different cell types, the cluster hypothesis provided in ***Demont-Guignard et al. (2012)*** was not adapted to NeoCoMM. Instead, the weakly synchronized firing phenomenon responsible for HFOs was obtained by simply increasing the NMDA conductance for all PCs (all layers). Thus, creating a depolarization with APs of a subgroup of PCs wherein the majority of cells did not fire. This supports the hypothesis of HFOs being mainly the result of enhanced excitation as opposed to tapered inhibitory transmission in the neocortex ***Lai et al. (2023)***. Another difference between the models was the fact that HFOs were generated without a depolarizing GABA (*E*_*GABA*_ = −75*mV*) as was the case for the hippocampus model. The percentage of the firing PC cells determined the nature of HFOs where higher firing cell percentage (between 20% and 45%) resulted in lower frequency band oscillations and lower depolarized PCs with APs gave way to FR (< 15%). This is consistent with other studies showing that pathological HFOs are the result of out-of-phase co-firing of small groups of interconnected and epileptic (hyperexcitable) PCs ***Zijlmans et al. (2012)***; ***de Curtis et al. (2012)***; ***Lai et al. (2023)***.

Besides its capability to realistically simulate various types of IEEs, the model also allowed us to study how postsynaptic current conductance values influence the transition between different IEE patterns. The results of this analysis reinforced the excitation-inhibition imbalance level principle discussed earlier for the generation of IES, SW, and DSW. In this respect, higher inhibitory postsynaptic currents were found to require a further increase in the excitatory conductances to achieve the desired event (Figure 5). In the case of HFOs, as explained in the previous part, their generation seems to be independent of the GABA postsynaptic current intensity. Based on this study, our model has allowed us to draw recommendations regarding electrophysiological parameters boundary values for the simulation of IEE patterns (Appendix 1-Table 1). A summary of the different pathophysiological parameters of IESs, SWs and HFOs generation mechanisms can be found in Table 1.

**Table 1.**
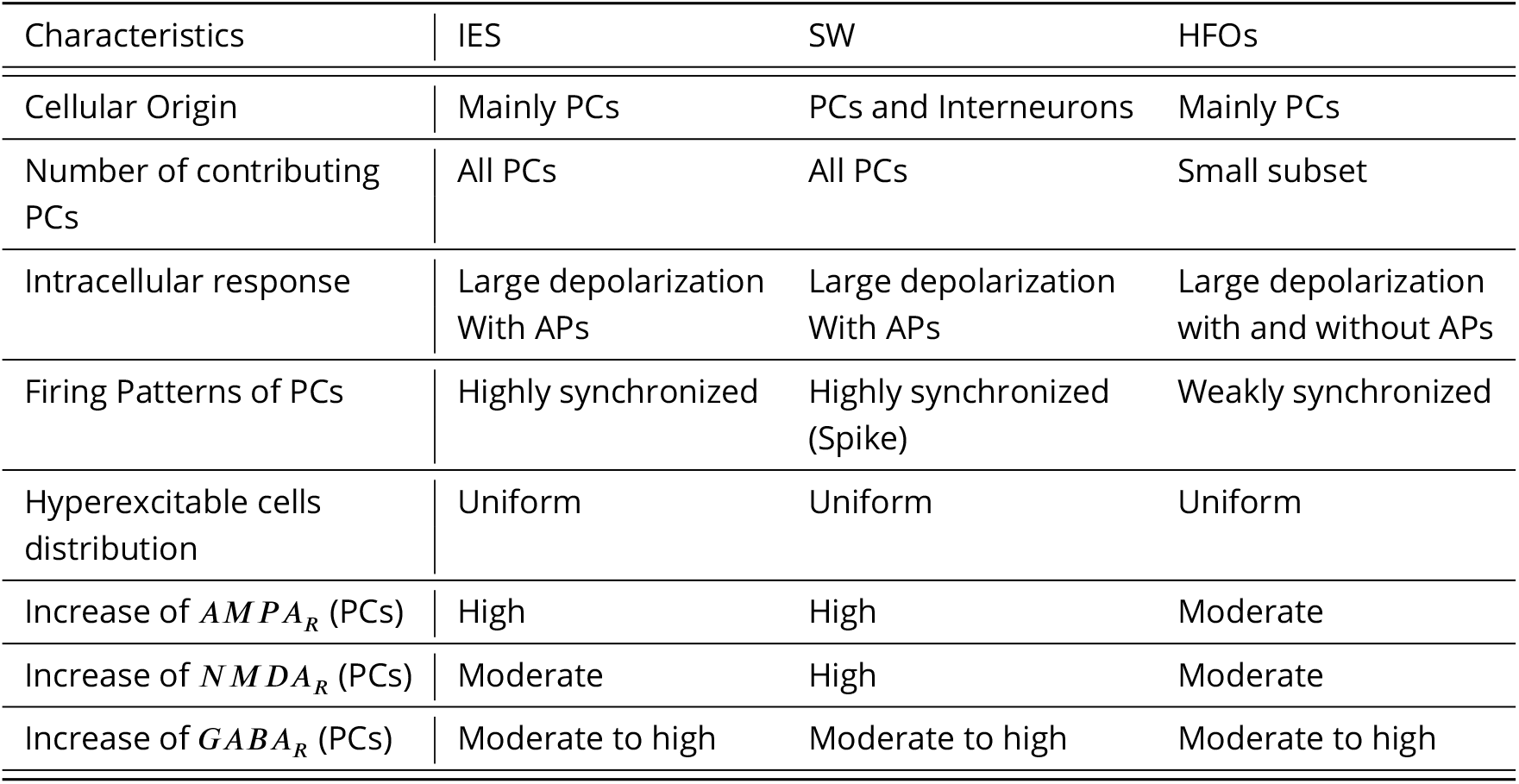
Summary of pathophysiological characteristics of different Interictal Epileptiform Events (IEEs). IES: Interictal epileptic Spike, SW: Spike and Wave, HFOs: High Frequency Oscillations, PCs: Principal Cells, *AMP A*_*R*_: *α*-amino-3-hydroxy-5-methylisoxazole-4-propionic acid receptor. *NMDA*_*R*_: N-methyl-D-aspartate receptor, *GABA*: Gamma-aminobutyric acid receptor.

| Characteristics | IES | SW | HFOs |
| --- | --- | --- | --- |
| Cellular Origin | Mainly PCs | PCs and Interneurons | Mainly PCs |
| Number of contributing PCs | All PCs | All PCs | Small subset |
| Intracellular response | Large depolarization With APs | Large depolarization With APs | Large depolarization with and without APs |
| Firing Patterns of PCs | Highly synchronized | Highly synchronized (Spike) | Weakly synchronized |
| Hyperexcitable cells distribution | Uniform | Uniform | Uniform |
| Increase of $AMPA_R$ (PCs) | High | High | Moderate |
| Increase of $NMDA_R$ (PCs) | Moderate | High | Moderate |
| Increase of $GABA_R$ (PCs) | Moderate to high | Moderate to high | Moderate to high |

Given the substantial differences in neural connectomics and anatomy between humans and rodents ***Loomba et al. (2022)***, we used our model to investigate the mechanisms and parameter settings required for the simulation of IEEs generated in the mouse’s cerebral cortex. Mainly, IEEs observed in the mouse model of epilepsy (Kainate mouse model of epilepsy) were used to validate our simulations including IESs, FRs, and repetitive spiking (Figure 12). Accordingly, we investigated the pathophysiological parameters responsible for the generation of these events using the mouse model settings in NeoCoMM. Interestingly, our results have highlighted the significant role of synaptic transmission parameters in interneurons for achieving the desired mechanisms responsible for IEEs generation. In fact, to simulate IEEs, it was necessary to increase not only the excitatory post-synaptic current of PCs but also those of the interneurons. This finding can be explained by the fact that, aside from the difference in neuron density and numbers, the locally projecting interneurons are lower for rodents compared to humans ***Vanderhaeghen and Polleux (2023)*** and the number of interneurons is 2.5 fold lower for mice compared to humans ***Loomba et al. (2022)***. In the case of HFOs simulation, the model allowed us to investigate the different subgroups (FR type 1 and type 2) of FRs that are usually observed in real recordings ***Frauscher et al. (2017)***. It highlighted the role of postsynaptic current conductance values in interneurons. Furthermore, in the case of FRs, it demonstrated that the same mechanism of weakly synchronized firing observed in human simulations can be replicated in mice by increasing the AMPA postsynaptic current of SST+ and VIP+ cells. However, this modification results in a simulation of Type 1 FRs, which are independent of spikes. To simulate Type 2 FRs, which occur simultaneously with spikes, it was necessary to reduce the firing of PV+ interneurons by decreasing *AMP A*_*R*_ and increasing *GABA*_*R*_ conductances.

Lastly, the presented model incorporated the biophysical model of the electrode contacts used in both clinical (SEEG electrodes) and in vivo recordings (twisted wire electrodes). This extension improved the realism of the model, as all the presented simulations included the ETI that integrates the actual geometrical and physical characteristics of the electrodes used in both human and mouse cases. By utilizing this feature, investigations into the geometrical and positioning characteristics of the SEEG electrode contacts enabled us to draw conclusions about the optimal radius, distance, depth, and orientation of the electrode contact to achieve the highest amplitude SWs. This tool could be employed for further analysis of optimized electrode designs for the recording of specific epileptic events in the future.

In this study, we introduced the NeoCoMM model as a new electrophysiologically reliable microscale computational model for simulating IEEs and investigating their underlying mechanisms. However, it offers a wide range of applications and can be highly beneficial in other analyses, such as assessing the impact of current stimulation, pharmacological modeling, designing electrodes for specific biomarkers, or conducting hypothesis testing. It provides an alternative to more complex, computationally demanding models while retaining essential physiological and biophysical aspects necessary for accuracy.

Limitations of the proposed model reside in the use of the same electrophysiological equations of voltage-dependent currents for all considered species. However, the connectivity, number of cells, density, and column morphology were adapted for each species. Moreover, adjusting other parameters allowed us to compensate for this shortfall. Another aspect that is lacking from the model is the neuroplasticity of the neural network. Still, in the case of the short simulations presented in this paper, plasticity is irrelevant. Future work will provide a newer version of the model which includes both physiological and electrophysiological plasticity. In addition, computing time will be further reduced using parallel computing which will allow us to propose the possibility of simulating multiple neocortical columns along with the interactions between them.

## Methods and Materials

### Neocortical Computational Microscale Model (NeoCoMM)

In this work, we propose a new physiologically realistic model of the cortical patch that incorporates its microcircuitry across the six layers. This model can be adapted to either Human, rats, or mice tissues and can be freely downloaded from https://gitlab.univ-rennes1.fr/myochum/neocomm.

#### Anatomical Structure of the Neocortical Column

The architecture of the neocortical column is defined by the type of cells it includes, their distribution in the column, the dimension of the different layers, and the connections between these cells.

#### Diversity of Modeled Neurons

The cells in the model were divided into two main classes: Principal Cells (PC), that are Glutamatergic excitatory neurons, and GABAergic inhibitory InterNeurons (IN). Based on their morphologies PCs were further divided into five types: Tufted Pyramidal cells (TTPC), Untufted Pyramidal cells (UTPC), Inverted Pyramidal Cells (IPC), Bipolar Pyramidal Cells (BPC) and Spiney Stellate Cells (SSC) ***Markram et al. (2015)***; ***Narayanan et al. (2016)*** (Figure 9.A). In total, PCs account for 70% to 80% of neurons in the neocortex and the rest are INs (Appendix 2-Table 1). The distribution of PC types across the layers is inhomogeneous and is portrayed in Appendix 2-Table 2. A 3D simplified representation of the PC cells’ main structural elements: soma, dendrites, and axon were defined for each type of cell (Appendix 2-figure 1). The dimensions of these volumes were adapted from ***Wang et al. (2018)*** for each layer and each species. Based on the neuromarkers they express, INs are comprised of four main types: the calcium-binding protein parvalbumin (PV+) expressing INs, the neuropeptides somatostatin (SST+) expressing INs, the vasoactive intestinal peptide (VIP+) expressing INs, and the protein reelin (RLN+) expressing INs ***Wamsley and Fishell (2017)***. In this study, each IN type is represented by one cell type except the PVs that are divided into Basket cells (BC) that inhibit the soma of PCs and Chandelier cells (ChC) that target the Axon Initial Segment (AIS) of PCs ***Wamsley and Fishell (2017)***. The other three types target the dendrites of other cells and are portrayed in this model by Martinotti Cells (MC) for SST, Bipolar Cells (BiC) for VIP, and Neurogliaform Cells (NGF) for RLN (Figure 9.A. Their distribution in the neocortical layers was adapted from ***Markram et al. (2015)*** and is depicted in 2-Table 2. Similarly to the PCs, the INs are represented by simplified 3D volumes wherein dimensions were obtained by averaging values from different studies (Appendix 2-figure 1) ***Wamsley and Fishell (2017)***; ***Laturnus et al. (2020)***; ***Niquille et al. (2018)***; ***Deleuze et al. (2019)***; ***Urban-Ciecko and Barth (2016)***; ***Prönneke et al. (2015)***; ***Cadwell et al. (2016)***.

**Figure 9.**
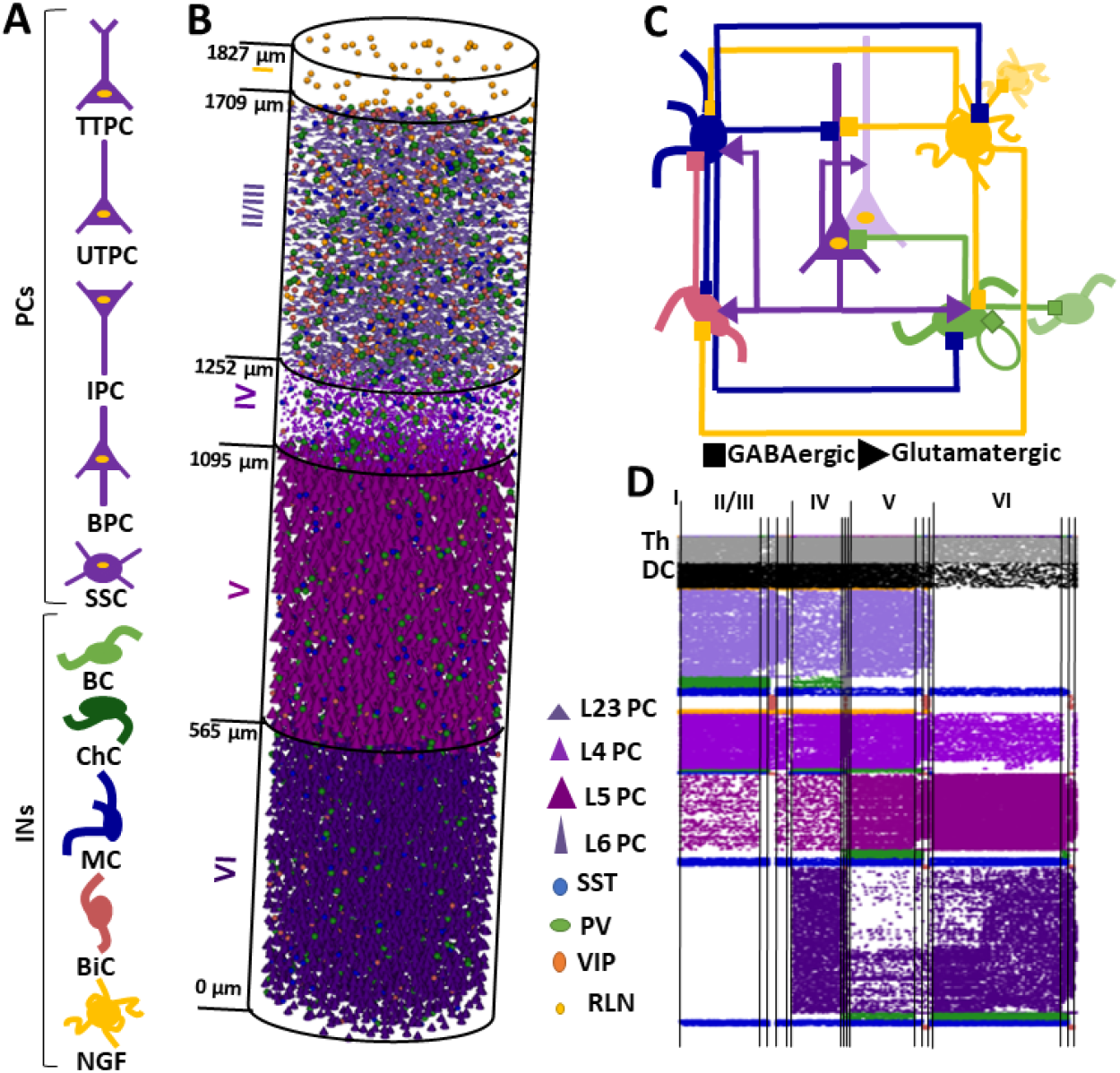
Anatomy of a cortical patch. A) The different cell types included in the model: Tufted Pyramidal cells (TTPC), Untufted Pyramidal cells (UTPC), Inverted Pyramidal Cells (IPC), Bipolar Pyramidal Cells (BPC) and Spiney Stellate Cells (SSC), Basket cells (BC), Chandelier cells (ChC), Martinotti Cells (MC), Bipolar Cells (BiC) and Neurogliaform Cells (NGF). B) A 3D rendering of the cortical patch with all the cell types for 13760 cells. C) The synaptic connectivity affinity diagram between the different cell types. D) an example of the connectivity matrix computed as described in section

**Table 2.**
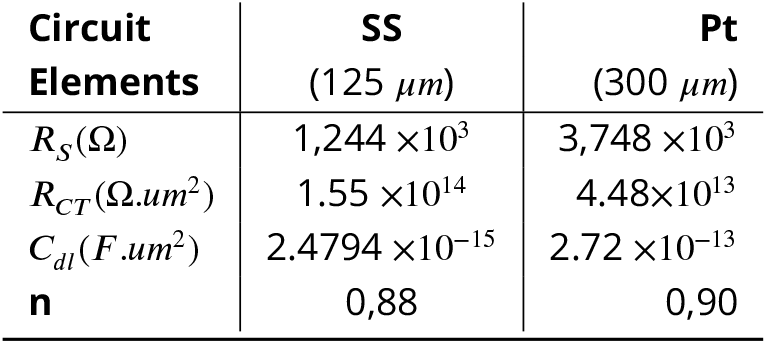
Values of the equivalent circuit elements used in the ETI model for the Platinum (Pt) and stainless steel (SS) electrode contacts.

| Circuit Elements | SS<br>(125 $\mu\text{m}$) | Pt<br>(300 $\mu\text{m}$) |
| --- | --- | --- |
| $R_s(\Omega)$ | $1,244 \times 10^3$ | $3,748 \times 10^3$ |
| $R_{CT}(\Omega.\mu\text{m}^2)$ | $1.55 \times 10^{14}$ | $4.48 \times 10^{13}$ |
| $C_{dl}(F.\mu\text{m}^2)$ | $2.4794 \times 10^{-15}$ | $2.72 \times 10^{-13}$ |
| n | 0,88 | 0,90 |

#### Cortical patch structure

Depending on the selected species, the 3D cortical patch structure was delineated as a cylinder with a radius of 210 *μm* and varying heights: 2622 *μm* for humans, 1827 *μm* for rats, and 1210 *μm* for mice, as detailed in prior studies ***Markram et al. (2015)***; ***DeFelipe et al. (2002)***; ***DeFelipe (2011)***.

This cortical patch is composed of six layers, wherein layers II and III were combined due to their similar characteristics. The thickness of each layer is different depending on the species and is defined following ***Defelipe et al. 2002 DeFelipe et al. (2002)***. The somas of different cells were placed inside each layer respecting their distribution using the best candidate algorithm ***Mitchell (1974)***. The number of cells in each layer was computed following the neuronal density measured in ***DeFelipe et al. (2002)***. An Example of the 3D neocortical volume simulated in this study is depicted in Figure 9.B. For this example, we modeled a human neocortical column of 1376 cells.

#### Connectivity of the Microcircuit

The synaptic connectivity of the microcircuit was determined following several steps. Based on Peter’s rule ***Braitenberg and Schüz (2013)***, connectivity between two cells is determined by the overlapping of neurites. Accordingly, Using the 3D volumes defined for each type and subtype of cells in each layer of the neocortical patch, we computed the overlap of each cell’s dendrites with all other cell axons ***Packer et al. (2013)***. For the connections between PVs (source) and PCs (target), the overlapping was computed between the axons of the PCs and the soma/AIS of the PV for BC/ChC respectively ***Wamsley and Fishell (2017)***; ***Deleuze et al. (2019)***. The connection between two cells respected several rules (see Figure 9.C):

- VIP cells only inhibit SST cells
- RLN cells do not receive intracortical excitatory input ***Jiang et al. (2013)***
- SST cells do not have autaptic connexions ***Laturnus et al. (2020)***
- PV cells inhibit only PCs and other PV cells in their own layers ***Deleuze et al. (2019)***

An afference matrix was also defined that outlines the percentage of afferences for each cell type following its type and layer ***Denoyer et al. (2020)***; ***Wamsley and Fishell (2017)***; ***Urban-Ciecko and Barth (2016)***; ***Wamsley and Fishell (2017)***; ***Jiang et al. (2013)***; ***Tremblay et al. (2016)***; ***Karnani and Jackson (2018)*** (Appendix 3-table 3). Respecting these rules, connectivity and weight vectors were determined for each cell describing the list of presynaptic cells and the corresponding synaptic weight. An example of the connectivity matrix is shown in Figure 9.D. The presynaptic weight was computed as the normalized 3D volumetric superposition of volumes of pre- and post-synaptic neurites ***Hill et al. (2012)***. The external input consisted of excitatory input from the Distant Cortex (DC) and Thalamus (Th) portrayed by Pyramidal cells from layers II/III (40 %) and V/VI (60 %). The external input number was defined as 7 % of the total number of PCs in the cortical patch ***Peters and Feldman (1976)***; ***Denoyer et al. (2020)***. These connections are added to the final connectivity matrix as shown in Figure 9.D.

#### Electrophysiological Models of Individual Cells

Electrical diversity of principal cells: The three compartments reduced model

A reduced conductance-based model of three compartments was used for the modeling of PCs. This model was adapted from the two-compartments model of Demont et al. ***Demont-Guignard et al. (2009)***. It consisted of three separate compartments (1) soma (2) dendrites and (3) AIS that were coupled via two conductances as portrayed in Figure 10. The membrane potential variation for each compartment was computed following the electric charge conservation equation described in the following differential equations:

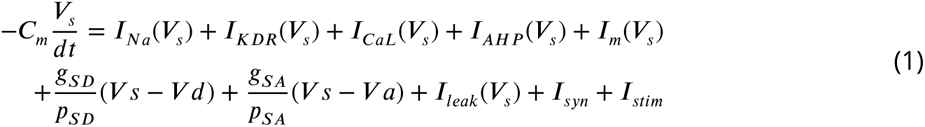

**Figure 10.**
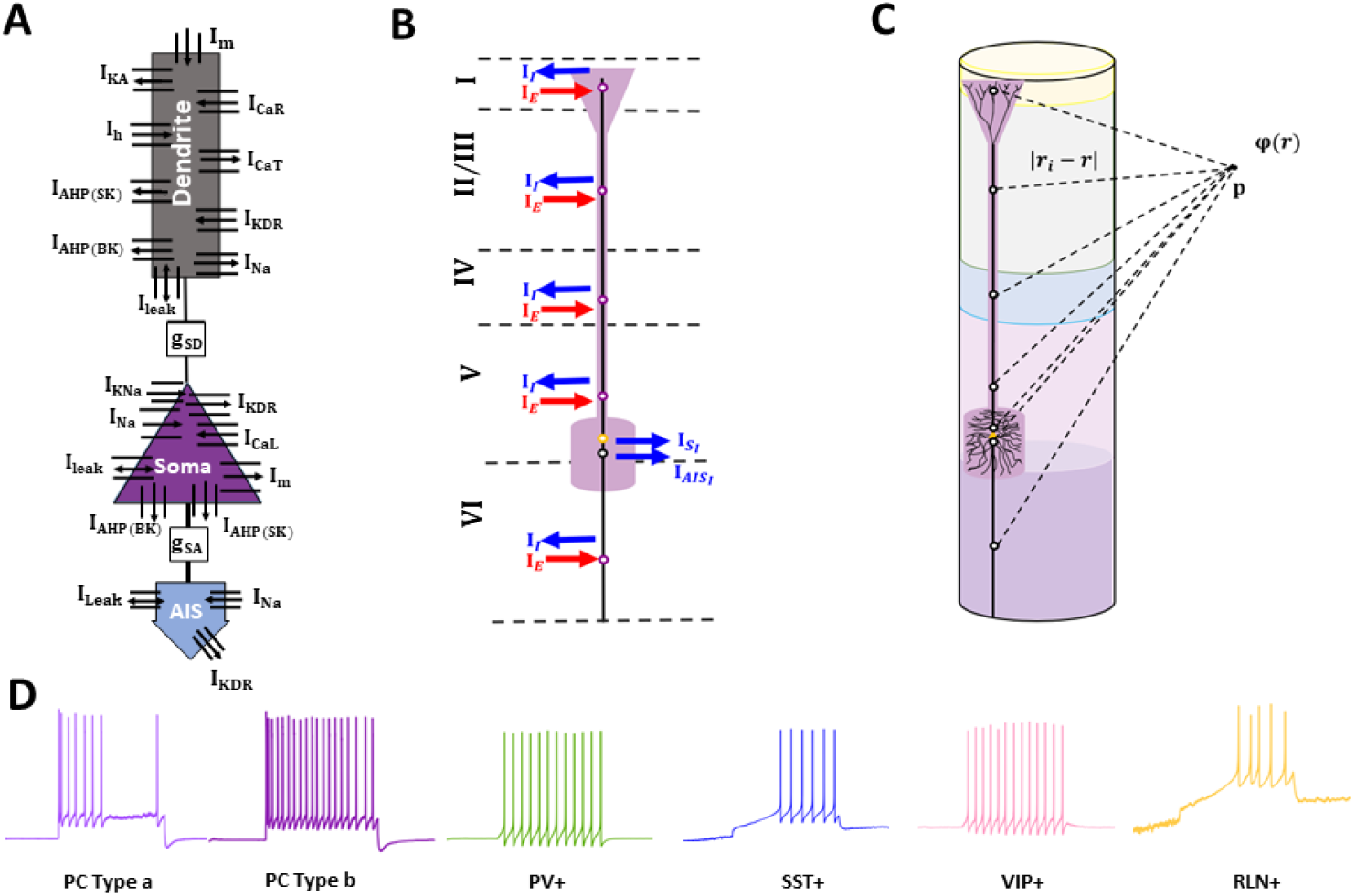
**A**.Pyramidal neuron computational model with three compartments: Soma, Dendrites, and Axon initial segment (AIS). All compartments have Voltage dependent sodium current (*I*_*Na*_), potassium delayed rectifier (*I*_*KDR*_) and a leak currents (*I*_*l*_ *eaK*). The soma has a muscarinic potassium current (*I*_*m*_), calcium-dependant potassium currents (*I*_*AHP*_), and an L-type calcium current (*I*_*C*_ *aL*). The dendrtite have *I*_*m*_, *I*_*AHP*_, T- and R-type calcium current (*I*_*C*_ *aT*, *I*_*C*_ *aR*), inactivating potassium current (*I*_*KA*_) and hyperpolarization-activated catonic current (*I*_ℎ_). coupling between compartments is acheived though conductances *g*_*SD*_ and *g*_*SA*_. **B**. Diverse firing patterns of the different cells modeled in the neocortical circuit in response to a depolarizing step current injected into the cell’s soma. **C**.A schematic of the Local Field Potential (LFP) reconstruction process.

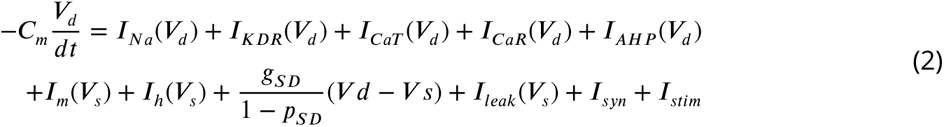

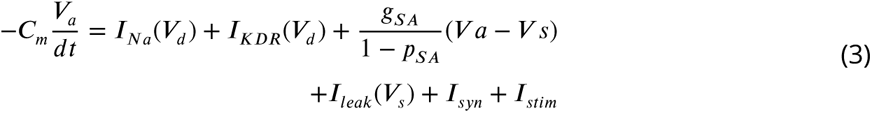

Where *V*_*s*_, *V*_*s*_ and *V*_*a*_ are the membrane potentials of the three compartments (soma, dendrites and AIS), *C*_*m*_ is the membrane capacitance, *g*_*SD*_ and *g*_*SA*_ are the conductances between soma/dendrites and soma/AIS respectively, *p*_*SD*_ and *p*_*SA*_ are the proportions of the soma area to the sum of soma/dendrites and soma/AIS respectively, *I*_*stim*_ is the external stimulation and *I*_*syn*_ is the sum of the synaptic currents. Each compartment had a different set of channels. The key active ionic currents chosen for the soma and dendrites accounted for seven and ten different voltage-gated channels respectively (Figure 10) ***Demont-Guignard et al. (2009)***. The AIS compartment had only two voltage-gated channels, the sodium *I*_*Na*_ and the Potassium delayed release *I*_*KDR*_ obtained from the Traub model ***Traub et al. (1994)***. All compartments had a leak current that portrayed the resting membrane potential variation with a Gaussian noise. The equations of the ionic currents followed the Hudgkin-Huxley formalism: *I*_*ion*_ = *g*_*ion*_*m*^*x*^ℎ^*y*^(*V*_*m*_ − *E*_*ion*_) equation with *g*_*ion*_ the ionic conductance, *x* and *y* are the number of gate activation and inactivation variables respectively, *E*_*ion*_ the reversal potential and *V*_*m*_ the membrane potential of the compartment. Gating variables dynamics are detailed in ***Demont-Guignard et al. (2009)*** for the soma and dendrites and in ***Traub et al. (1994)*** for the AIS. Passive properties were set as: 1 *μC* ∕*cm*^2^ for the soma and AIS membrane capacitance (*C*_*m*_) and 2 *μF* ∕*cm*^2^ for the dendrites, 0.18 *mS*∕*cm*^2^ for the mean leak conductance (*g*_*l*_*eaK*) and -70 *mV* for the resting potential (*E*_*r*_), 1 *mS*∕*cm*^2^ for *g*_*SA*_ and *g*_*SD*_. In the context of neocortical pyramidal cells, the conductance values of ion channels for the three compartments were adapted in order to portray a firing rate profile similar to the one recorded from pyramidal cells in the neocortex ***Zhang et al. (2017)***; ***Mitrić et al. (2019)***. Accordingly, we defined two main groups of electric types for the PCs: layers II/III and IV and layers V and VI. The values of conductances used in this model are presented in Appendix 4 Table 1.

#### Electrophysiological Model of Interneurons

The interneuron models consisted of one a compartment model ***Hajós et al. (2004)*** with various numbers of voltage-gated channels that were adapted to portray the four different types of firing patterns of interneurons used in the model (PV+, SST+, VIP+, and RLN+). An example of the firing pattern of each cell type in response to a depolarizing step current injected into the soma is presented in Figure 10.D

#### Synaptic Diversity and External input

For excitatory synaptic connections, both *α*-amino-3-hydroxy-5-methyl-isoxazole-4-propionic acid (AMPA) and N-methyl-D-aspartic acid (NMDA) receptors (*AMP A*_*R*_, *NMDA*_*R*_) were modeled. The corresponding glutamatergic postsynaptic currents were obtained following ***Hajós et al. (2004)***; ***Destexhe et al. (1994)***. Similarly, GABA(ergic) synaptic currents were modeled based on ***Hajós et al. (2004)***. A constant (*ω*) for each synaptic connection was added to these equations in order to adjust the weight of the excitatory and inhibitory connections received by each cell. The (*ω*) value is obtained by normalizing the corresponding synaptic input weight vector in the weight matrix obtained from the connectivity reconstruction algorithm described in section *Connectivity of the Microcircuit*.

#### Reconstruction of LFP

The Local Field Potentials (LFP) is considered to be the signal resulting from extracellular electrical potentials around the recording electrode ***Einevoll et al. (2013)***. In the cortex, these potentials derive from the transmembrane currents of principal cells wherein synaptic input is the primary contributor. Accordingly, the LFP at a recording point *p* was computed based on the biophysical forward modeling volume conductor theory. The extracellular medium is presumed to be infinite, isotropic, and homogeneous. Currents entering and leaving the cell compartments are considered sources and sinks respectively. For each three-compartment principal cell *i*, its contribution to the extracellular potential (Φ) is obtained by the sum of the individual potentials evoked by the presynaptic currents and their accompanying return currents at the recording point *p* as expressed by equation (4) ***Nunez and Srinivasan (2006)***.

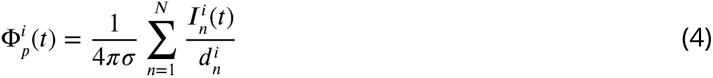

With 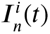 the *n*^*t*ℎ^ transmembrane current, 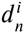 the distance between the recording point and 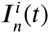 locations, *N* is the total number of transmembrane currents and *σ* the extracellular conductivity. The positions of the transmembrane currents were determined based on the biophysical propagation of excitatory/inhibitory synaptic currents across the multi-compartments of a neuron. A schematic illustration is depicted in Figure 10.C.

The total potential received by the active surface area of the electrode can be calculated by integrating the previously defined extracellular potential equation across the entire surface area. This is approximated as follows ***Fuglevand et al. (1992)***:

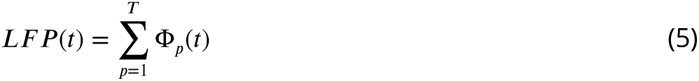

with *T* representing the total number of discretization points. The step size used for discretization is set at 10 *μm*.

#### Electrode Tissue Interface (ETI)

A classical Randles Model was used as an equivalent circuit to simulate the recording electrode employed in SEEG recordings. This model comprised spreading resistance (*R*_*s*_) in series with the double layer which includes a charge transfer resistance (*R*_*CT*_) in parallel with a pseudo-capacitance (*Z*_*CP A*_) (Figure 7). For more information about the ETI, please refer to ***Al Harrach et al. (2023)***. The LFP recorded by the electrode is obtained after computing the inverse Fourier transform of the product between the extracellular potential V(f) and the ETI transfer function H(f). The circuit element values for the SEEG electrode’s cylindrical contact of 300 *μm* diameter and 2 *mm* height and the wire disc contact of 125 *μm* are given in Table 2. These values were adapted from ***Al Harrach et al. (2023)***; ***Franks et al. (2005)***. The appropriate electrode transfer function H(f) was applied for all the simulations presented in this study for human and mouse settings.

### SEEG Data

The interictal clinical SEEG signals used in this study were obtained from recordings that took place at the Epilepsy Surgery Department of La Timone University Hospital in Marseille, France. They are part of a larger database of signals collected after authorization from the Institutional Review Board (IRB00003888, IORG0003254, FWA00005831) of the French Research Institute of Health and Medical Research (Inserm). SEEG signals were recorded from patients during presurgical evaluation after informed consent and being aware of their potential use for research purposes. SEEG electrodes placement was personalized based on medical information related to the epileptogenic zone and controlled using telemetric X-ray imaging. A Deltamed-NatusTM system with 256 channels equal to 256 was used for the recording. The sampling frequency was set to 1024 Hz (to verify). A hardware analog high-pass filter (cut-off frequency equal to 0.16 Hz) was present in the recording system to remove very slow oscillations of the baseline. For this particular study, SEEG signals were chosen in a patient with electrode contacts in the neocortical regions. Figure 11 presents examples of the patient’s X-rays showing different implanted SEEG along with segments of the recorded signals. An example of SW, IES, and HFOs is also illustrated in Figure 11.C.

**Figure 11.**
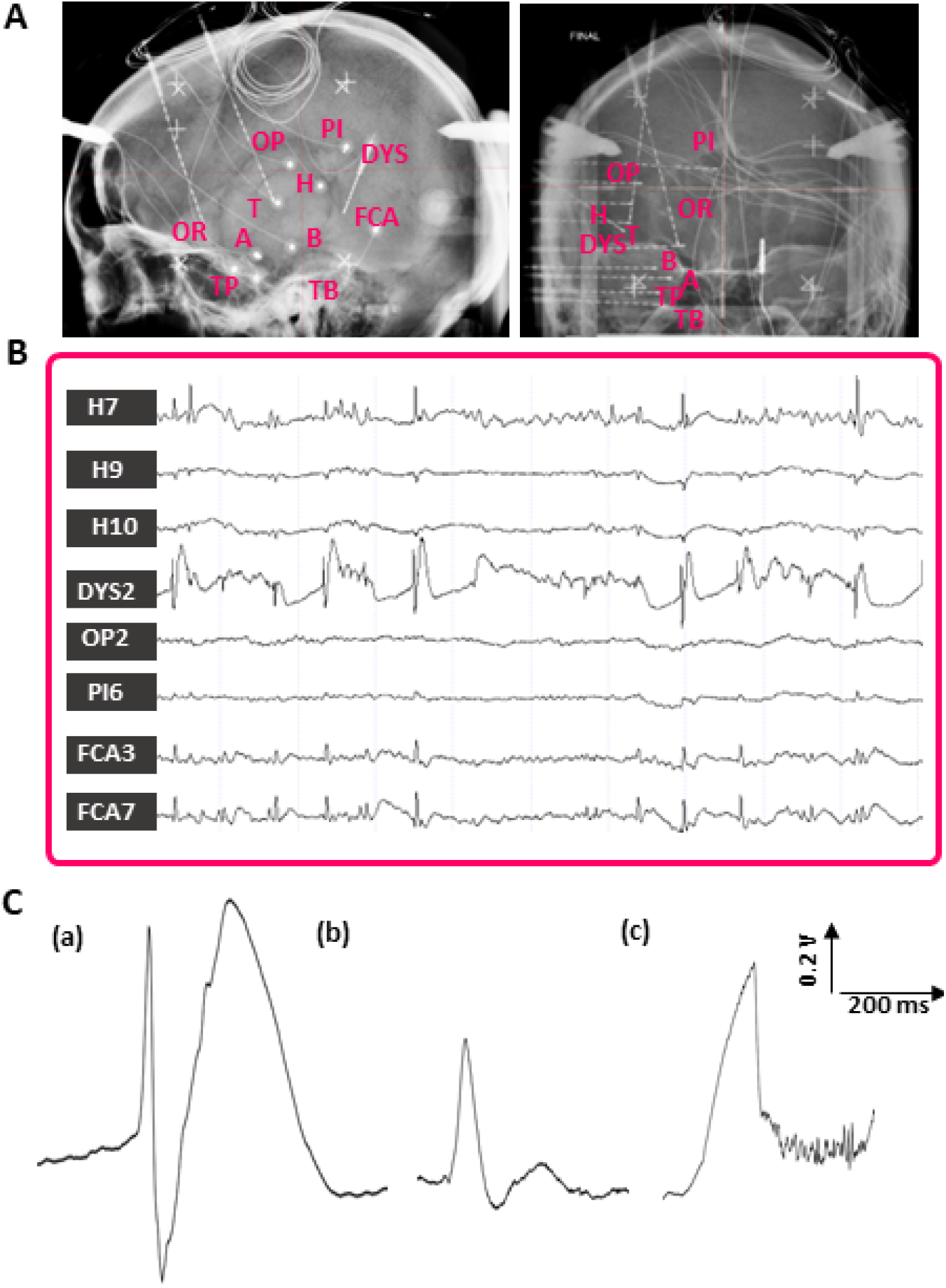
Interictal Epileptic Events (IEEs) recorded with intracranial electrodes from a human epileptic cortex. A) Skull X-rays (sagittal and coronal) of a patient illustrating the SEEG electrodes positioning. B) example of monopolar interictal SEEG signals. The visualized signals are a subset of a larger set of 128 channels. The electrodes are positioned as follows: TP: Temporal Pole, TB: Temporal Basalis, FCA: Fissura Calcarina Anterior, H: Heschl Gyrus, OR: frontal orbital, OP: parietal operculum, PI: Sub parietalis, FCA: anterior calcarine sulcus, T: anterior T1, A: amygdala, B: anterior Hippocampus. C) examples of interictal Spike and wave (SW) (a), Spike (b), and Ripples (c).

### Animal model

For the in vivo experimental validation of the Model, we used the signal recorded from epileptic mice following the iron chloride mouse model of epilepsy ***Jo et al. (2014)***. This experiment respected the European Communities Council Directive of 24 November 1986 (86/609/EEC) and was approved by the ethics committee on animal experimentation in Rennes, France (agreement 23603). The electrode implantation was performed three months after *F eCl*_3_ injection. The mice were placed in a stereotaxic device during implantation (Figure 12.A). Four stainless steel (SS) electrodes were inserted into the somatosensory cortex through drilled burr holes (Figure 12.B). Among these electrodes, two were of 125 *μm* diameter and were placed at AP= -0.5 mm, ML= +1.5 and -1.5 mm, DV= 0.7 mm (coordinates from Bregma). The other two were of 250 *μm* diameter and were inserted at AP = +0.5 and -3.5 mm, ML = -1.5 mm, DV= 0.5 mm. A reference electrode of 125 *μm* was placed on the bone. Surgical glue and dental cement were used to fix the electrode. In this work we only considered the recordings from the 125 *μm* radius microelectrode in the ipsilateral hemisphere. An example of a 25 *s* recording is shown in Figure 12.B wherein IEE segments were highlighted and are depicted in Figure 12.C for IES, HFOs, and repetitive spiking.

**Figure 12.**
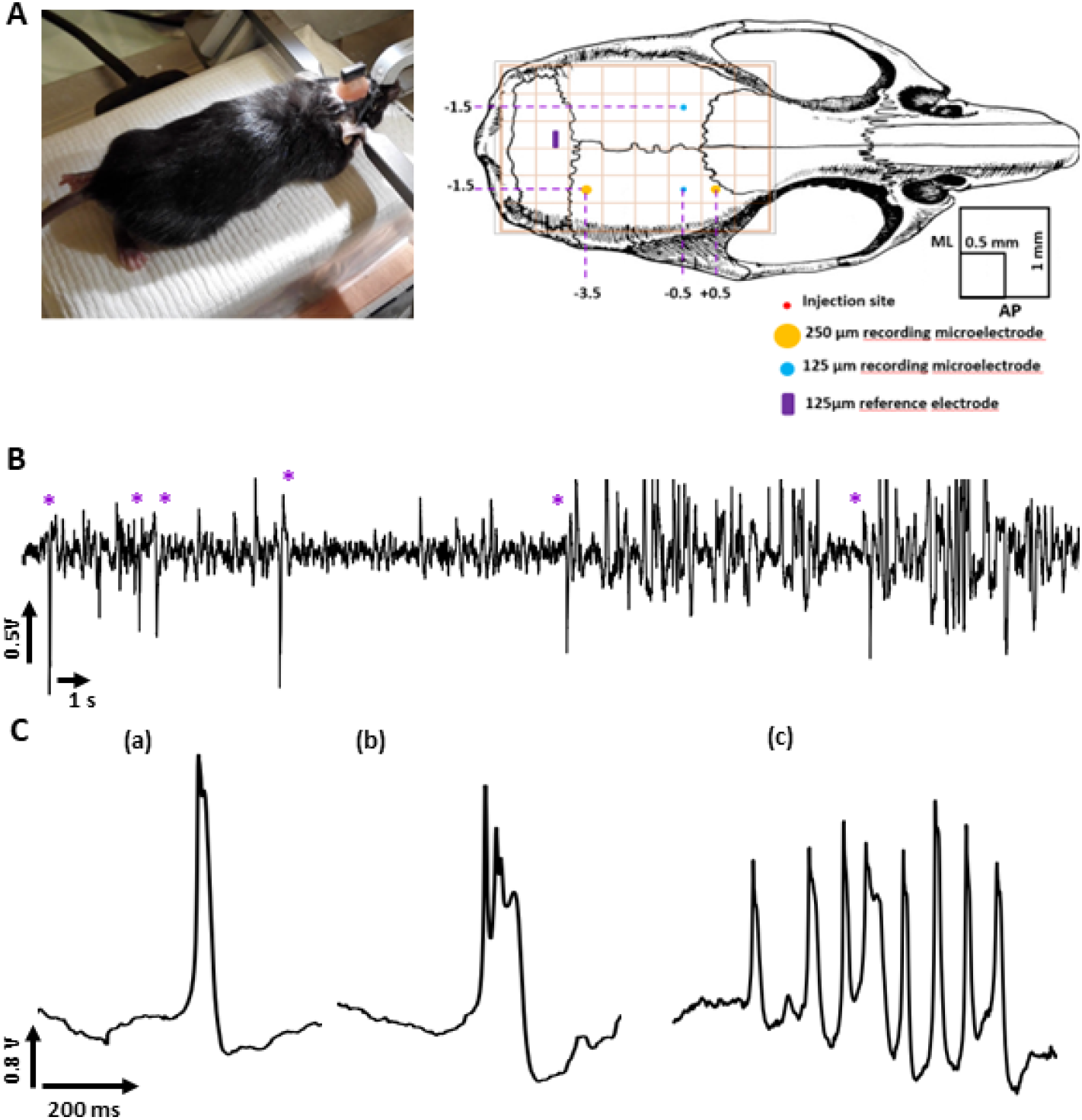
Interictal Epileptiform Events (IEEs) in iron chloride epilepsy model. (A) Image of the operating field during electrode implantation (the mouse is fixed in a stereotaxic frame) with a schematic diagram of the multisite intracortical electrode implantation positions. AP: antero-posterior, ML: mesio-lateral. (B) A segment of the recorded signal using the 125 *mum* radius microelectrode. The IEEs in the segments were depicted with an asterisk(C) Examples of interictal Spike (IES) (a), FR (b), and repetitive spiking (c).

## Acknowledgments

We would like to thank our Lab technician Mr Gabriel Dieuset (Laboratoire Traitement du Signal et de l’Image (LTSI-U1099), Université de Rennes 1, INSERM, 35000 Rennes, France) for the experimental data acquisition and his continuous help and support in the conception and application of experimental protocols. We would also like to thank the research engineer in the Galvani ERC Project Dr. Audrey Barbedette for her help in the in vivo recordings. This project has received funding from the European Research Council (ERC) under the European Union’s Horizon 2020 research and innovation program (grant agreementNo 855109)

## Appendix 1

**Appendix 1—table 1.** Electrophysiological parameter values for the simulation of different neocortical Interictal Epileptiform Events (IEE) in the Human neocortex with NeoCoMM.

|  | Spikes | Spike and Wave | Ripples | Fast Ripples | Double Spike |
| --- | --- | --- | --- | --- | --- |
| $E_{GABA}(mV)$ | [-75, -56] | [-75, -56] | [-75, -60] | [-75, -60] | [-75, -65] |
| $g_{GABA}(mS/cm^2)$ | [25, 40] | [25, 40] | [25, 40] | [25, 40] | [25, 35] |
| $g_{AMPA}(mS/cm^2)$ | [8, 12] | [8, 12] | [6, 8] | [6, 8] | [10, 12] |
| $g_{NMDA}(mS/cm^2)$ | [0.2, 0.6] | [0.4, 0.8] | [0.2, 0.8] | [0.2, 0.8] | [0.2, 0.8] |
| $Jitter(ms)$ | [4, 10] | [6, 10] | [6, 10] | [6, 10] | [8, 12] |

## Appendix 2

### Morphological characteristics of neocortical individual cells

**Appendix 2—table 1.**
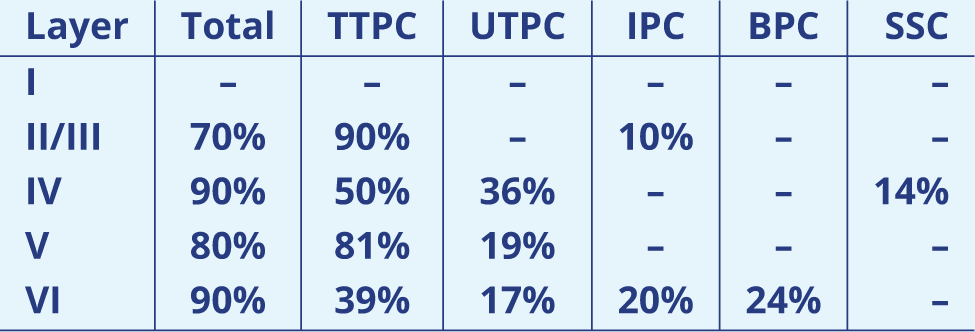
Distribution of the Principal cells (PCs) across the Layers of the neocortex ***Markram et al. (2015)***. TTPC: Tufted Pyramidal cells, UTPC:Untufted Pyramidal cells, IPC: Inverted Pyramidal Cells, BPC:Bipolar Pyramidal Cells and SSC: Spiney Stellate Cells.

**Appendix 2—table 2.**
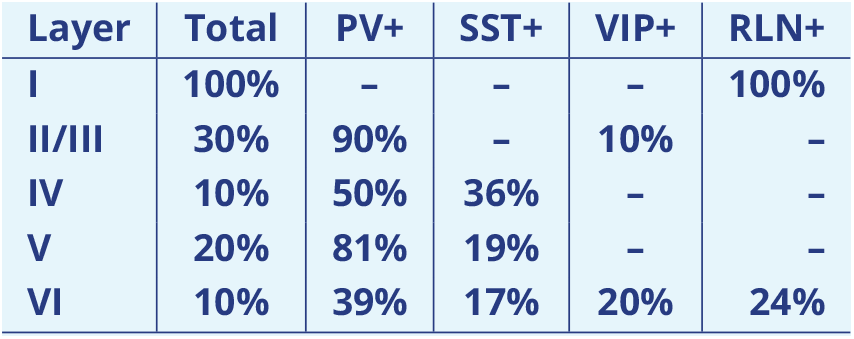
Distribution of the Principal cells across the Layers of the neocortex. PV+: parvalbumin, SST+: the neuropeptides somatostatin, VIP+: vasoactive intestinal peptide, RLN+: reelin.

| Layer | Total | PV+ | SST+ | VIP+ | RLN+ |
| --- | --- | --- | --- | --- | --- |
| I | 100% | - | - | - | 100% |
| II/III | 30% | 90% | - | 10% | - |
| IV | 10% | 50% | 36% | - | - |
| V | 20% | 81% | 19% | - | - |
| VI | 10% | 39% | 17% | 20% | 24% |

**Appendix 2—figure 1.**
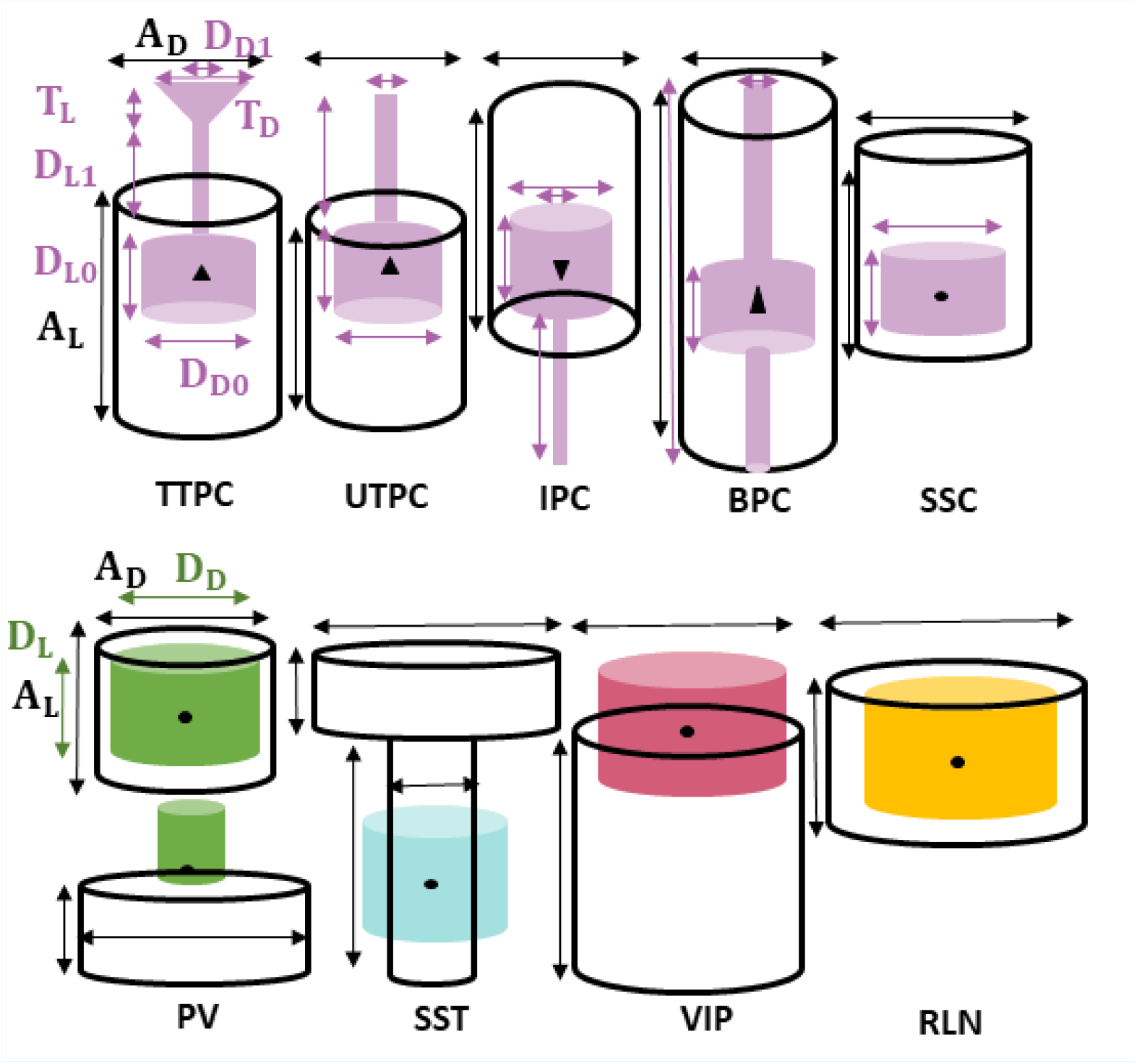
The 3D simplified representation of the different cells described in the neocortex. TTPC: TufTed Principal Cells, UTPC:UnTufted Principal cells, IPC: Inverted Principal Cells, BPC: Bipolar Principal Cells, SSC: Spiney Stellate Cells, PV: Parvubalmin expressing interneurons (divided into Basket cells (BAS, up) and Chandelier Cells (ChC, bottom)), SST: SomatoStatin expressing Cells, and VIP: Vasoactive Intestinal Peptide expressing cells. The dimension parameters values are depicted in

**Appendix 2—table 3.** Values of the 3D geometrical representation of the neurons soma, dendrites and axons depicted in Appendix 2-figure 1. The values are given in *μm*.

|  | II/III | IV | V | VI |
| --- | --- | --- | --- | --- |
| $S_d$ | 18 | 11.77 | 24.44 | 17.30 |
| $S_h$ | 6.32 | 7.57 | 23.90 | 16.94 |
| $A_d$ | 410 | 400 | 400 | 400 |
| $A_L$ | 735 | 1057 | 1014 | 682 |
| $D_{D0}$ | 206 | 272 | 293 | 243 |
| $D_{L0}$ | 164 | 144 | 150 | 139 |
| $D_{D1}$ | 10 | 10 | 10 | 10 |
| $D_{L1}$ | 25 | 216 | 329 | 114 |
| $D_{D2}$ | 195 | 186 | 293 | 286 |
| $D_{D2}$ | 275.8 | 499 | 792 | 593 |

**Appendix 2—table 4.** Values of the 3D geometrical representation of the interneurons soma, dendrites, and axons depicted in Appendix 2-figure 1. The values are given in *μm*.

|  | PV |  | SST | VIP | RLN |  |
| --- | --- | --- | --- | --- | --- | --- |
|  | BAS | ChC |  |  | I | II/III |
| $A_D$ | 200 | 300 | 150,450 | 150 | 400 | 120 |
| $A_L$ | 300 | 250 | 250,125 | 600 | 150 | 250 |
| $D_D$ | 150 | 50 | 100 | 60 | 150 | 150 |
| $D_L$ | 150 | 150 | 250 | 150 | 150 | 150 |

## Appendix 3

### Synaptic connectivity Computing

**Appendix 3—table 1.**
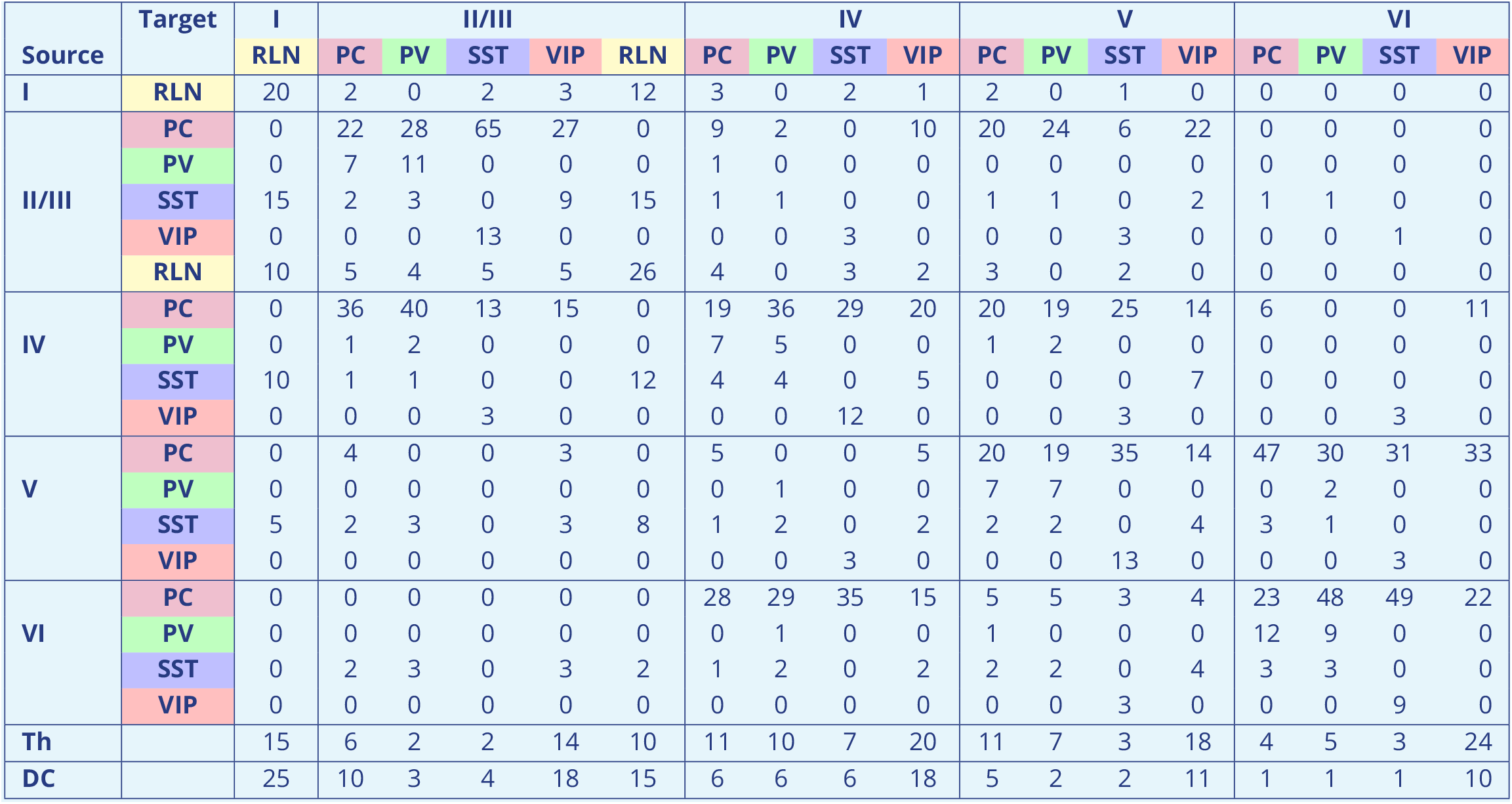
Synaptic afferent matrix. The percentage of afferent input for each cell type per layer in the neocortical column. PC: Principal Cells, PV: Parvalbumin expressing interneurons, SST: Somatostatin expressing interneurons, VIP: vasoactive intestinal peptide expressing interneurons, RLN: Reelin expressing interneurons, Th: Thalamocortical PCs, DC: distant cortex PCs. Values where adapted from several studies ***Urban-Ciecko and Barth (2016)***; ***Deleuze et al. (2019)***; ***Wamsley and Fishell (2017)***; ***Jiang et al. (2013)***; ***Karnani and Jackson (2018)***; ***Tremblay et al. (2016)***; ***Denoyer et al. (2020)***

## Appendix 4

### Electrophysiological parameters of neocortical individual cells

**Appendix 4—table 1.** ion channel Conductance values for the voltage-dependent currents defined in ***Demont-Guignard et al. (2009)*** for different cell types in *mS*∕*cm*^2^. *I*_*Na*_: Voltage-dependent sodium current,*I*_*KDR*_: potassium delayed rectifier,*I*_*m*_: muscarinic potassium current, *I*_*AHP*_: calcium-dependant potassium currents (**I*_*AHPSK*_*, **I*_*AHPBK*_*), **I*_*leaK*_*: leak, **I*_*CaL*_*: L-type calcium current, The T- and R-type calcium current (*I*_*CaT*_, *I*_*CaR*_), *I*_*KA*_: inactivating potassium current and *I*_*KNA*_: Sodium dependent potassium current.

|  | II/III/IV |  |  | V/VI |  |  |
| --- | --- | --- | --- | --- | --- | --- |
|  | Soma | dendrites | AIS | Soma | dendrites | AIS |
| $g_{Na}$ | 70 | 14 | 200 | 70 | 014 | 200 |
| $g_{KDR}$ | 6 | 2 | 200 | 6 | 2 | 200 |
| $g_m$ | 3.1 | 0.1 | – | 3.5 | 0.1 | – |
| $g_{AHP_{SK}}$ | 0.1 | 10 | – | 0.1 | 20 | – |
| $g_{AHP_{BK}}$ | 2 | 1 | – | 2 | 1 | – |
| $g_{CaL}$ | 0.5 | – | – | 0.5 | – | – |
| $g_{CaT}$ | – | 1 | – | – | 1 | – |
| $g_{CaR}$ | – | 3 | – | – | 0.75 | – |
| $g_h$ | – | 0.4 | – | – | 0.4 | – |
| $g_{KA}$ | – | 55 | – | – | 55 | – |
| $g_{KNA}$ | 0 | – | – | 0.5 | – | – |

**Figure 2—figure supplement 1.**
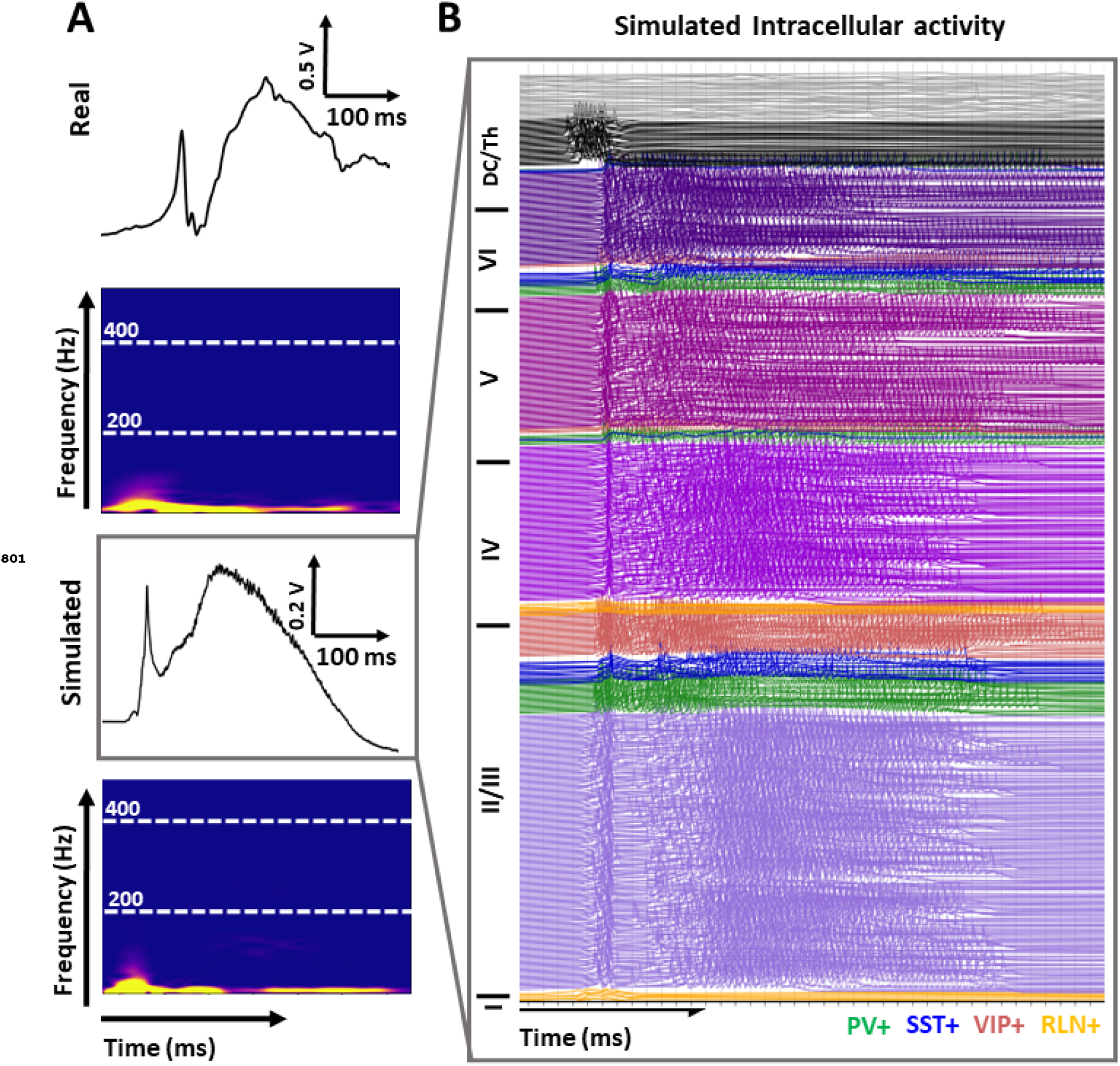
The intracellular activity corresponding to the SW signal in (A)

**Figure 2—figure supplement 2.**
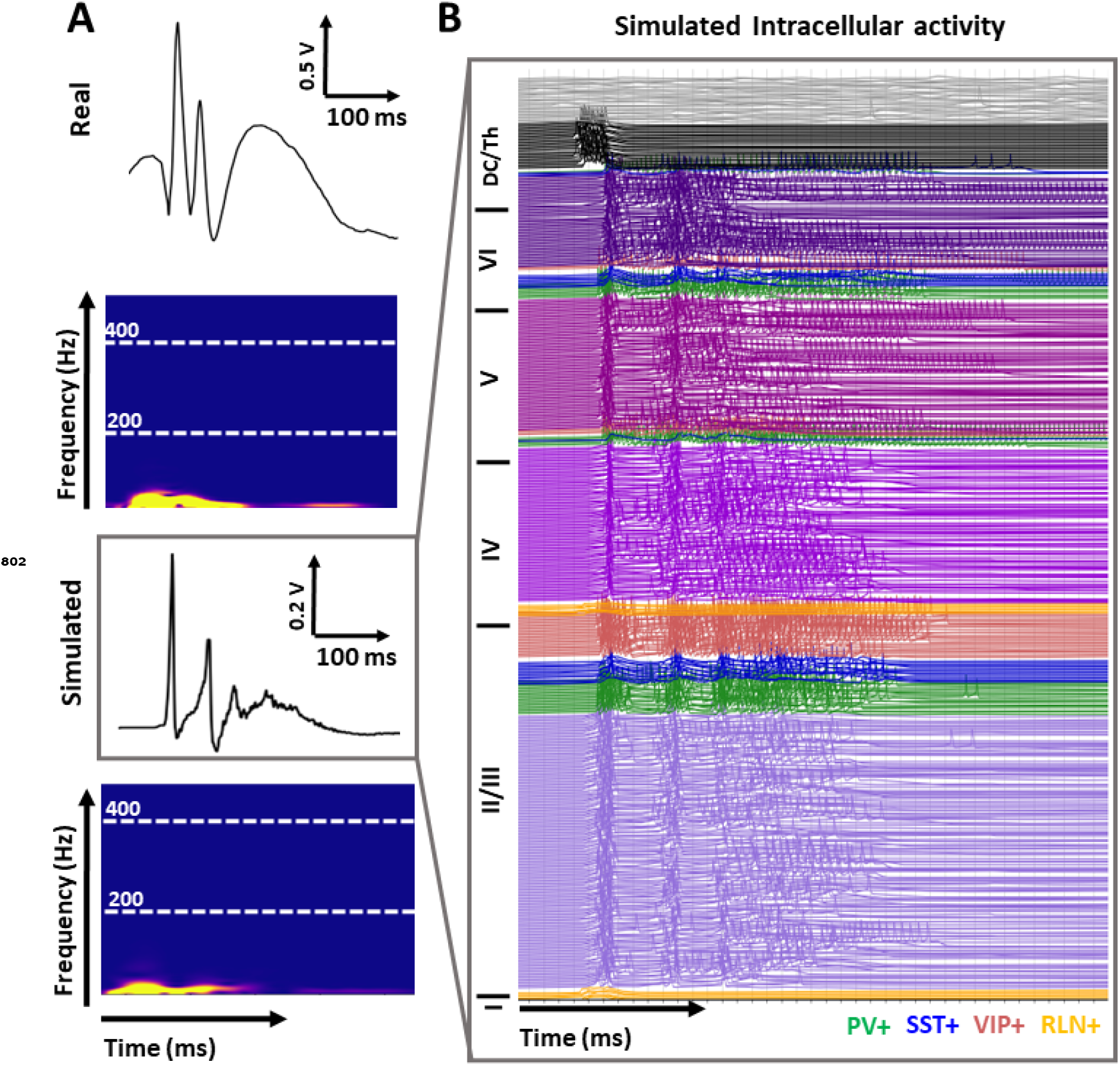
The intracellular activity corresponding to the DSW signal in (B)

**Figure 2—figure supplement 3.**
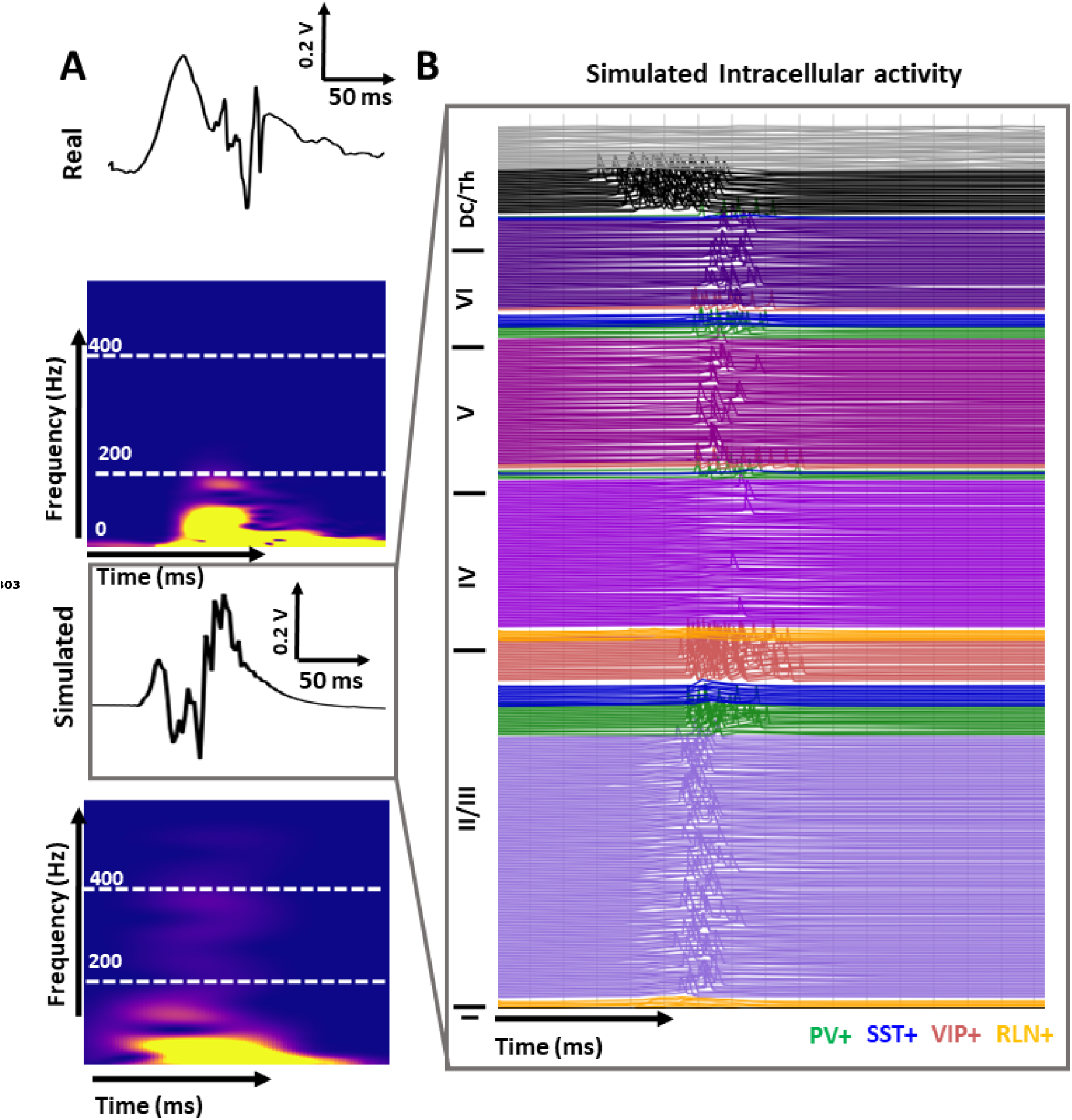
The intracellular activity corresponding to the HFOs signal in (C)

**Figure 2—figure supplement 4.**
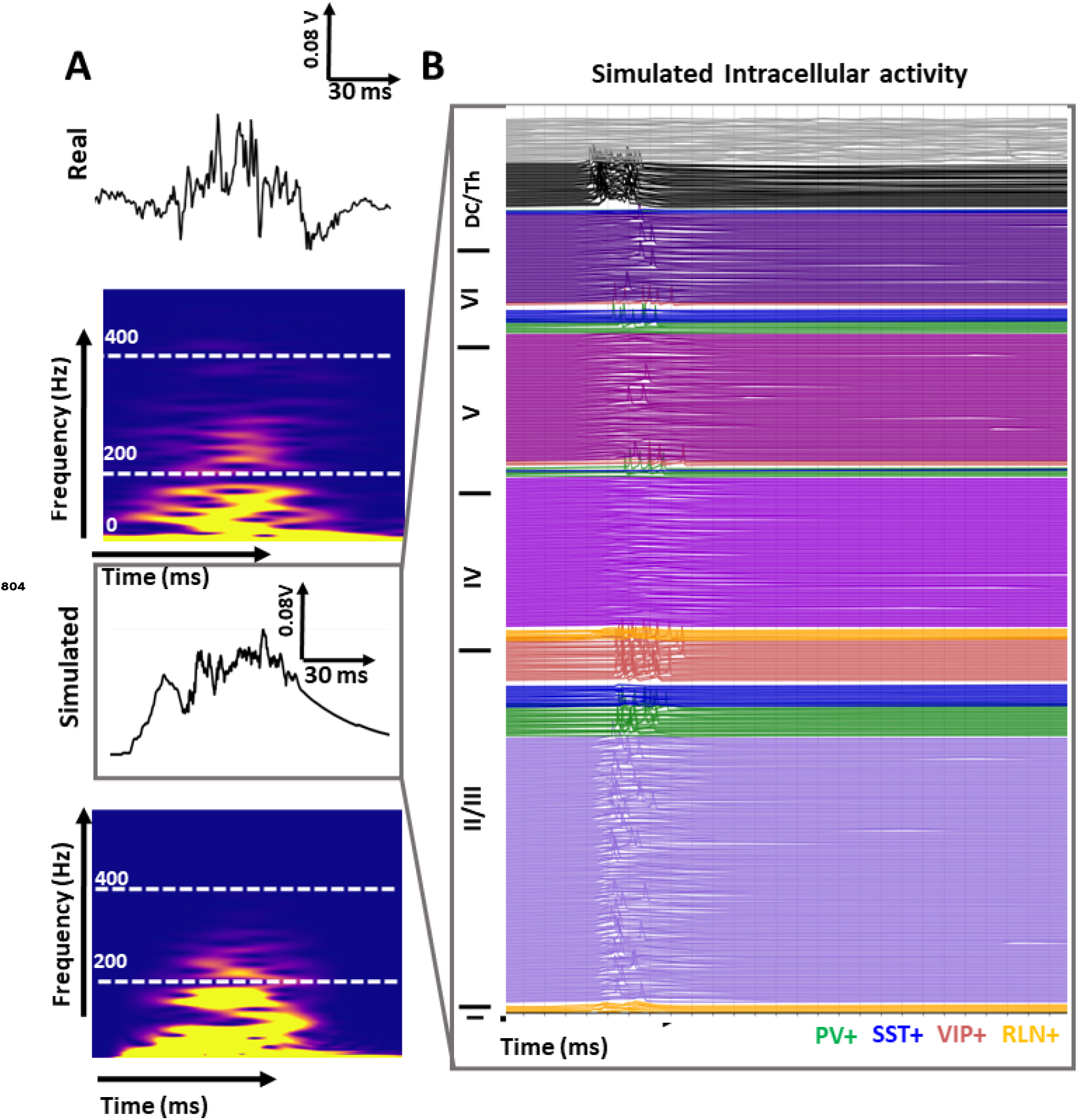
The intracellular activity corresponding to the FR signal in (D)

**Figure 6—figure supplement 1.**
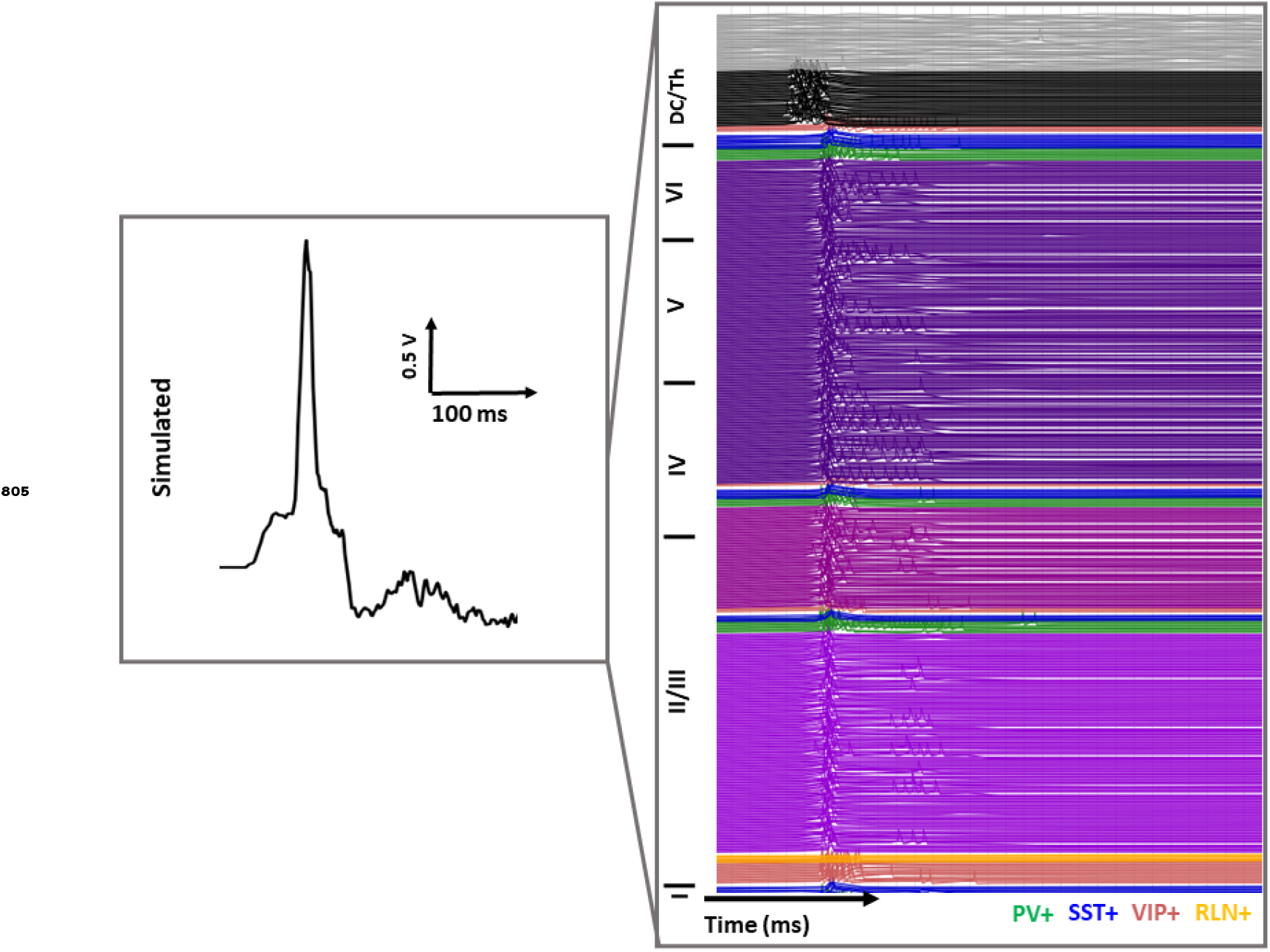
The intracellular activity corresponding to the IES signal in (A)

**Figure 6—figure supplement 2.**
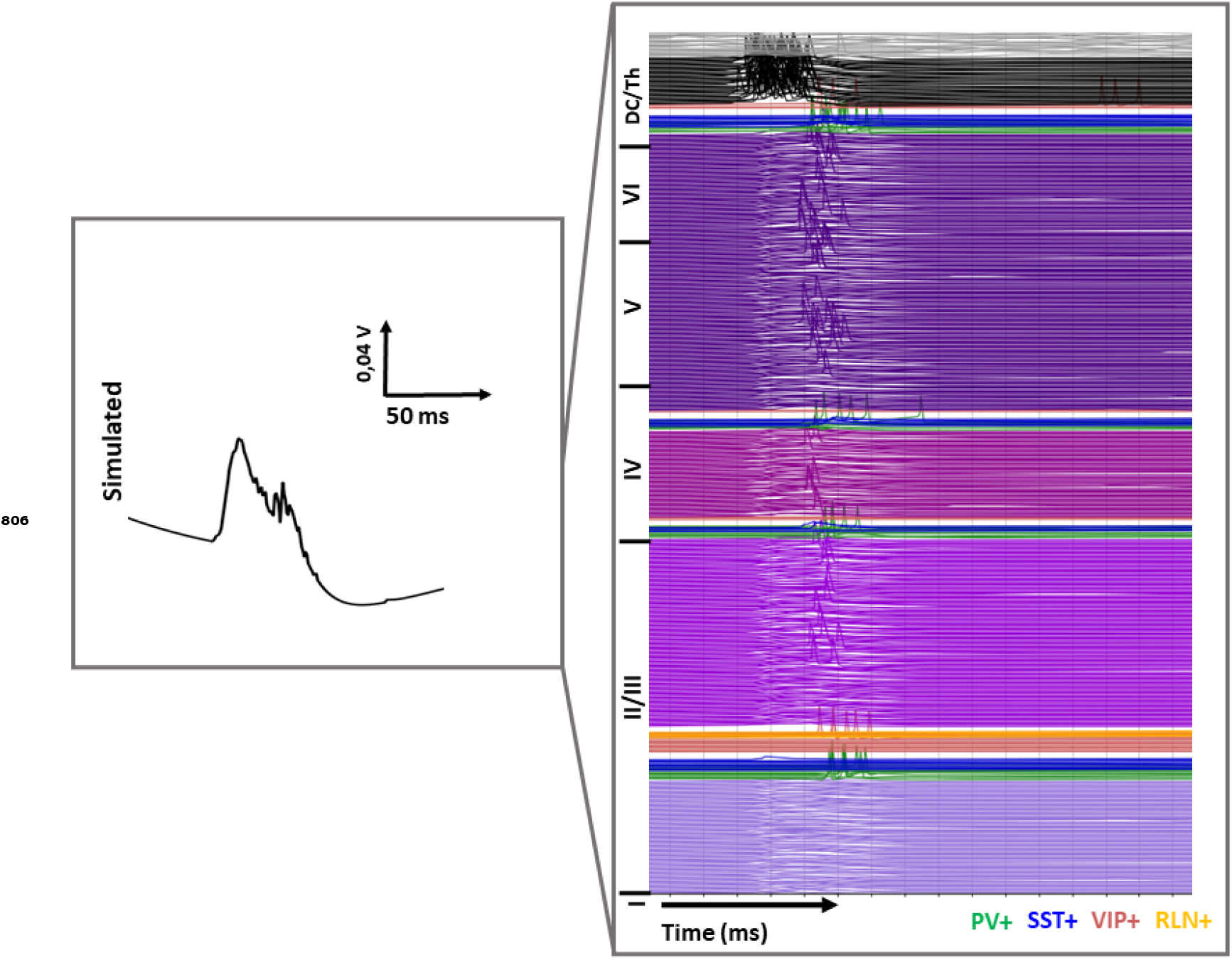
The intracellular activity corresponding to the FR type 1 signal in (B)

**Figure 6—figure supplement 3.**
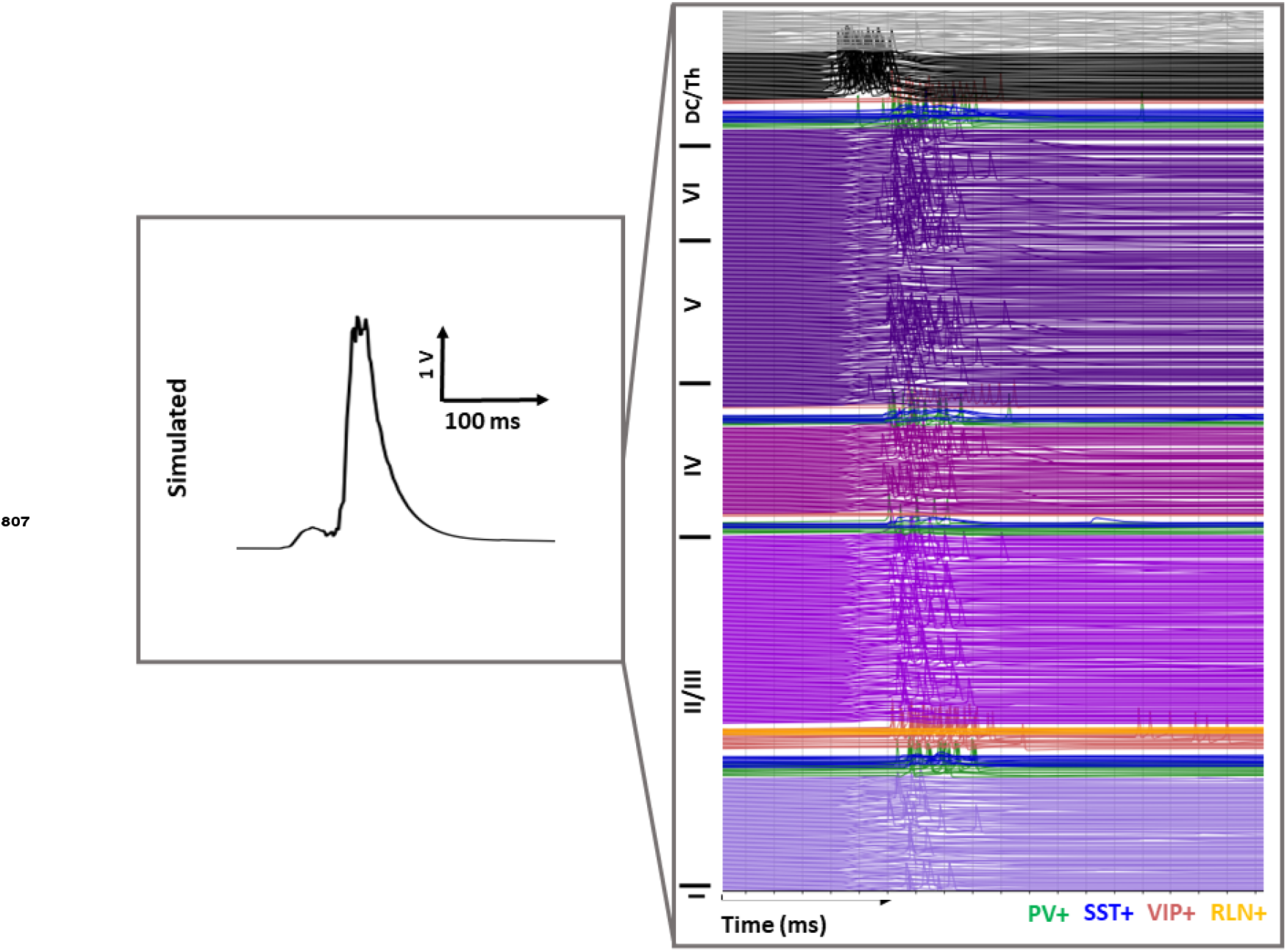
The intracellular activity corresponding to the FR type 2 signal in (B)

**Figure 7—figure supplement 1.**
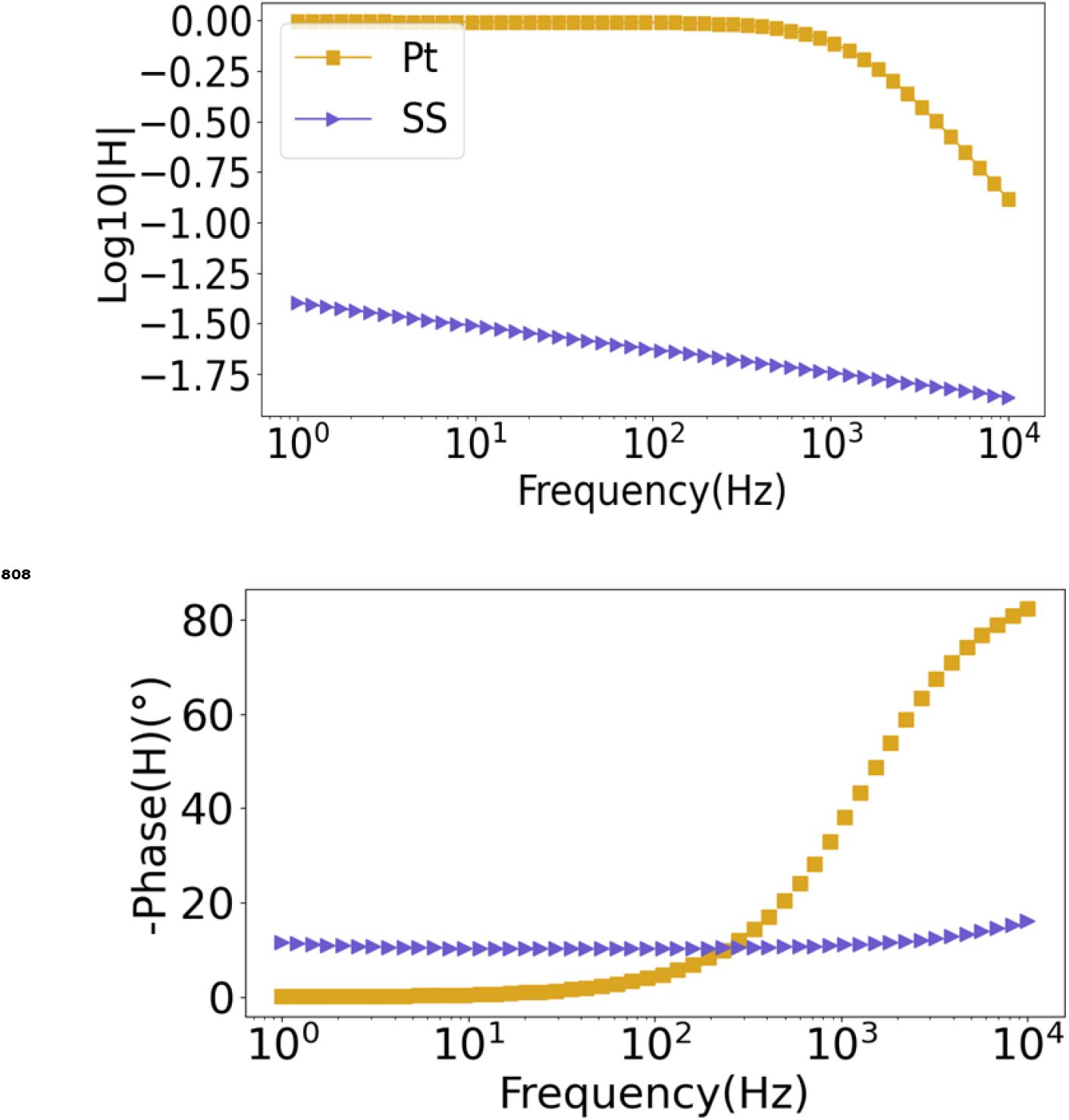
Bode plot of the transfer functions for both Pt and SS electrodes

